# A *Pan*-pangenome illuminates complex structural variation and selection in humans, chimpanzees, and bonobos

**DOI:** 10.64898/2026.06.06.730619

**Authors:** Joana L. Rocha, Runyang Nicolas Lou, Carolina Lima Adam, Prajna Hebbar, Scott Ferguson, Davide Bolognini, Alison Killilea, Kendra Hoekzema, Andrea Guarracino, Yun Deng, Nicole Soranzo, Benedict Paten, Erik Garrison, Alex Pollen, Evan E. Eichler, Rori V. Rohlfs, Matthew W. Mitchell, Peter H. Sudmant

## Abstract

Complete, haplotype-resolved genome assemblies have provided unprecedented insight into the evolution of structurally complex, rapidly evolving regions of human genomes^1–6^; however, population-scale pangenome resources of our closest relatives, chimpanzees and bonobos (genus, *Pan*), are necessary to ascertain the origins and evolutionary context of these loci. Here, we sequence and assemble 58 haplotypes from four distinct *Pan* clades to high contiguity (median contig NG50=54 Mb), including eight near-T2T genomes. These genomes reveal previously intractable genetic variation increasing estimates of genome-wide genetic diversity 6-37% across populations compared to short-read estimates^7^. We identify recurrent structural polymorphisms across species impacting genes associated with immune response and host-pathogen interaction and find that structural variants (SVs) are 170- to 260-fold more likely than single nucleotide variants (SNVs) to exhibit high-impact effects across species. Contrasting SV patterns across primates we find that transposable element mutation rates differ by as much as threefold between species. We show that human disease-associated short tandem repeat (TR) loci have uniquely expanded in humans sensitizing our species to these TR-expansion disorders. Physically phased haplotypes enable reconstruction of genome-wide genealogical histories, uncovering ancient, functional genetic variation maintained by balancing selection, as well as signatures of recent adaptation in chimpanzee subspecies. Several malaria-associated loci exhibit ancient structural polymorphism, including the African great ape–specific glycophorin (*GYP*) gene expansion. We characterize the sequence, structure, and composition of diverse glycophorin haplotypes in humans and chimpanzees. We identify independent malaria-protective *GYPA-B* fusion events in humans and novel chimpanzee glycophorin genes resulting from both ancient and recent fusion events demonstrating parallel adaptations to pathogen resistance across hominins. Together, our resource highlights the critical importance of nonhuman primate population-scale pangenomics for understanding the evolution of complex genome structures and the biodiversity of our endangered closest living relatives.

## Introduction

Long-read sequencing and haplotype-resolved genome assemblies have transformed human medical and evolutionary genetics. Extending upon the first telomere-to-telomore (T2T) complete human genome^8^ these fully phased and assembled genomes have enabled the study of complex loci involved in human disease and adaptation that are inaccessible to short-read sequencing^1–6^. Yet, the origins, ancestral state, and evolutionary context of these complex regions of the human genome cannot be fully understood without high-quality comparative genomics resources nor can genomic features unique to the human population be defined without population-based analyses of our most closest related species. The recent sequencing of T2T genomes from apes provides an essential starting point for such analyses^9,10^. However, long-read-based population-scale pangenomes of nonhuman primates do not yet exist.

Chimpanzees (*Pan troglodytes*) and bonobos (*Pan paniscus*) are sister taxa to humans and are >98% identical across the alignable genome despite ∼6 million years of divergence^9,11^. These species comprise a geographically structured group of populations with bonobos diverging from chimpanzees ∼1 million years ago (Mya) and individual chimpanzee subspecies splitting from one another ∼160-580 thousand years ago (kya)^7,12^ **(Fig. 1A).** Chimpanzees are keystone species with important roles in maintaining ecosystem health across diverse habitats. However, anthropogenic forces such as poaching and habitat loss have forced these charismatic taxa into endangered status^13,14^. The importance of chimpanzees and bonobos for understanding human evolution, adaptation, and behavior has long been appreciated with key contributions from scientific giants such as Jane Goodall^15^, Mary-Claire King, and Alan Wilson^16^. Chimpanzees, like humans, have complex social systems and behaviors and maintain diverse cultural traditions^17^ which may have evolved as a result of local adaptation to a wide range of habitats^18^. Furthermore, humans and chimpanzees exhibit susceptibility to similar pathogens including lentiviruses such HIV/SIV^19^, ebola-causing filoviruses^20^, anthrax^21^, malaria-causing plasmodia^22^, and leprosy-causing mycobacteria^23^. Chimpanzees are also susceptible to many of the same developmental and age-associated diseases as humans^24,25^ providing a key point of reference given our shared genetics. Evolutionary anthropologists have long acknowledged that studying chimpanzees and bonobos provides a critical lens to understand what makes us human. However, we have until now lacked the high-quality whole genome resources necessary to deeply interrogate the entirety of their genomes and our shared evolutionary past.

**Figure 1.**
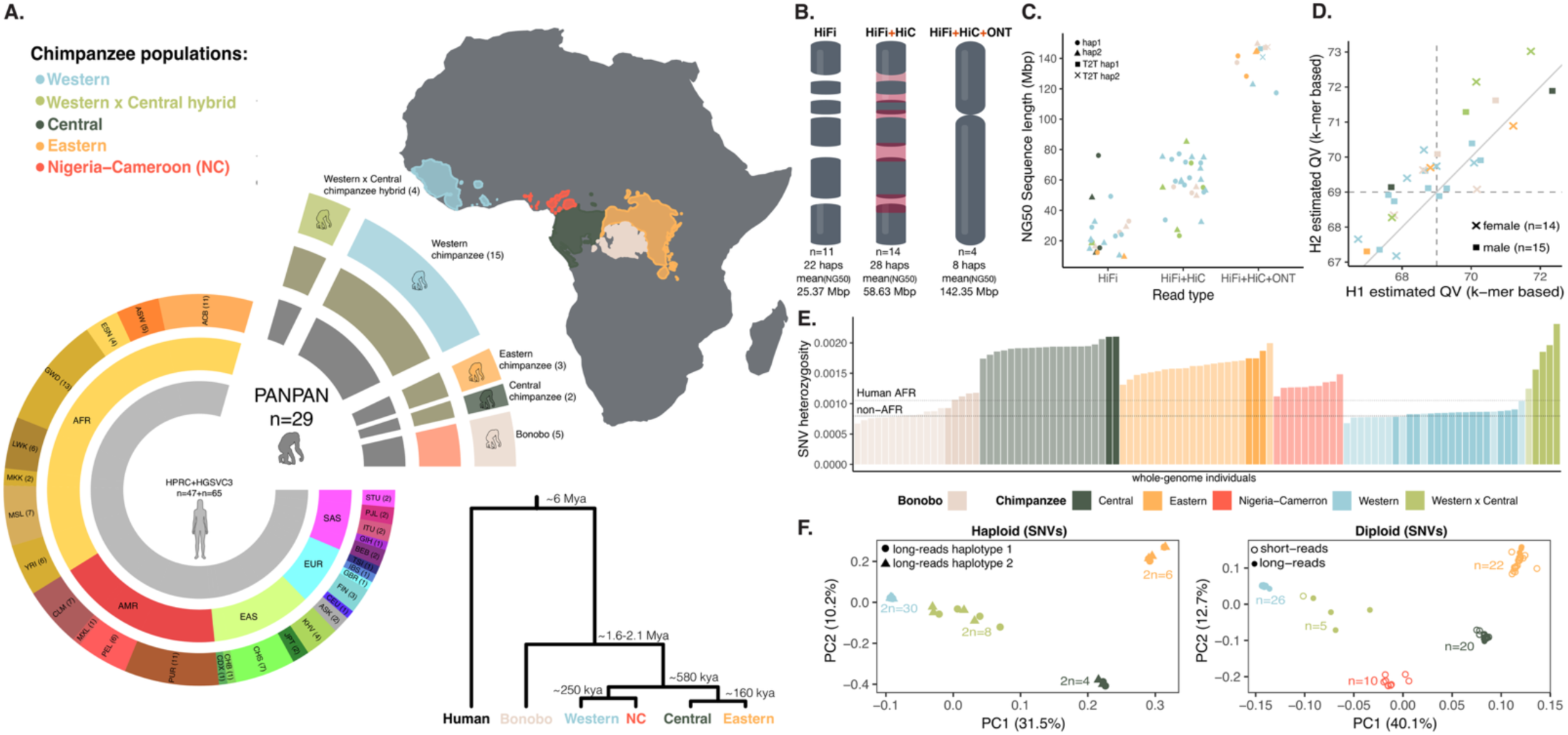
Population-scale sequencing and assembly of 58 diverse chimpanzee and bonobo haplotypes. **(A)** Overview of the 29 individuals (58 haplotypes) comprising this resource in the context of population-scale pangenome resources for diverse humans (HPRC and HGSCVC3). The cohort includes 24 chimpanzees (*Pan troglodytes*) representing three subspecies (Western, Central, Eastern) and Western x Central hybrids, alongside 5 bonobos (*Pan paniscus*). The map indicates IUCN geographic distribution alongside the phylogenetic relationship and divergence estimates (in kya - thousands of years ago, and Mya - millions of years ago) between populations and species of the genus *Pan* and humans. **(B)** Distribution of assembly methods used: PacBio HiFi only (n=11), PacBio HiFi + Hi-C (n=14), and HiFi + Hi-C + ONT (n=4, near-T2T). **(C)** Contiguity metrics for the 58 haplotypes. Assemblies generated with HiFi/HiFi+HiC reached an average NG50 of 44 Mb, while Verkko assemblies reached an average NG50 of 142.35 Mb, comparable to existing T2T references. **(D)** Quality Values (QV) for the assemblies, calculated via k-mer analysis. Values range from 67 to 73, indicating a base-level accuracy exceeding 99.9999%. **(E)** Genome-wide per-individual SNV heterozygosity across *Pan* species and populations, ordered by population and within-population diversity. Bar transparency distinguishes long read-sequenced individuals (n = 29, opaque) from Illumina short read-sequenced individuals (n = 72, semi-transparent). Dashed lines show average SNV heterozygosity for human HGDP–1KGP cohorts (AFR ≈ 1.05 × 10⁻³, n = 609; non-AFR ≈ 7.96 × 10⁻⁴, n = 2,024). **(F)** Principal component analysis (PCA) of chimpanzee SNV diversity mapped to the T2T mPanTro3 reference. Left: haploid PCA of SNVs called from long-read haplotype-resolved assemblies (n = 24 individuals, 48 haplotypes). Right: diploid PCA of SNVs jointly called from read-mapped data combining long reads from this study with short reads (n=59) from the Great Ape Genetic Diversity Project (total of 83 individuals total. Points coloured by population; shapes indicate data type. Axis labels give the variance explained by PC1 and PC2.

While the recent assembly of the complete genomes of six ape species unlocked previously intractable repetitive and complex sequences to evolutionary analysis^9^, individual reference genomes only begin to scratch the surface of the extraordinary diversity and complexity of structural haplotypes present across populations of different species^4,26,27^. Here we sought to construct a pangenome resource of 58 haplotypes from four distinct clades of diverse chimpanzees and bonobos using long-read approaches, drawing on both captive and wild-born males and females with available low passage cell lines (**Supplementary Table S1, Fig. 1A-B**). By integrating our resource with existing human haplotype assemblies, we characterize the structure, composition, functional importance, and evolutionary trajectories of genomic variation that is both unique to and shared between chimpanzees, bonobos and humans. Together, we demonstrate the critical importance of nonhuman primate population-scale pangenomics for understanding the evolution of genome structure.

## Results

### A population-scale resource of 58 diverse chimpanzee and bonobo assemblies including 8 near-T2T genomes

We utilized lymphoblastoid and fibroblast cells from 24 chimpanzees and 5 bonobos including representative individuals from three of the four recognized chimpanzee subspecies: Western chimpanzees (*P.t. verus, n=15*), Central chimpanzees (*P.t. troglodytes, n=2*), and Eastern chimpanzees (*P.t. schweinfurthii, n=3*), alongside n=4 Western x Central hybrid individuals (**Supplementary Table S1, Fig. 1A**). For all samples we extracted high molecular weight DNA and performed long-read Pacific Biosciences (PacBio) high-fidelity (HiFi) sequencing to an average of 50x (**EDFig. 1A).** Across a subset of 18 individuals we additionally generated Hi-C sequencing data to enable long-range genome scaffolding and phasing (HiFi+HiC genomes) and for 4 of these we further generated Oxford Nanopore Technologies (ONT) Ultra Long Read sequencing (∼55.5x) to enable near-T2T genome assembly (HiFi+HiC+ONT, **Fig. 1B**). We constructed haplotype-resolved genome assemblies for all individuals with HiFi-asm^28^ and assembled the 4 HiFi+HiC+ONT genomes using Verkko^29^.

The resulting set of 58 haplotype assemblies are geographically representative, highly contiguous and base-pair accurate, complementing and extending existing long-read haplotype resolved T2T genomes (**Fig 1B-D, EDFig. 1A-F**). Average contig NG50 values of ∼44 Mb were achieved for the the 50 HiFi and HiFi+HiC haplotype assemblies (**Fig. 1B-C**, **EDFig. 1B-C**) while the NG50 of the 8 HiFi+HiC+ONT assemblies reached ∼142.35 Mb (auNG 143.3Mb, **Fig. 1B-C**, **EDFig. 1B-C**). These 8 haplotypes thus approach the T2T quality of the recent complete ape genomes mPantro3 and mPanPan1 which exhibit NG50s of 143.56Mb and 147.25Mb, respectively. However, we note that only 23.4% of the chromosomes among these 8 near-T2T haplotype assemblies were complete (gapless with a telomere on both ends; **EDFig. 1E-F, Table S2-S4**) in contrast to 74% of the chromosomes in ape T2T reference genome assemblies. All 58 haplotype-resolved genomes had assembly quality values (QV) between 67 and 73 (**Fig. 1D**) similar to the level of T2T primate genomes and average single copy BUSCO scores of ∼97.5% (**EDFig. 1D**). Together, these genomes represent the largest base-level accurate, haplotype-resolved resource of nonhuman primate population diversity data to date.

Previous studies aiming to catalogue chimpanzee and bonobo genetic variation using short-read whole-genome sequencing have provided key insights into diversity patterns and population structure^7,12,30,31^. Still, short-reads cannot accurately capture complex regions of the genome, leaving critical within-species variation uncharacterized. To illustrate how long-read sequencing data can complement and be integrated with previously generated whole-genome sequencing datasets we jointly called 47,107,694 biallelic single nucleotide variants from long- (n=24) and short-read (n=59) chimpanzee data mapped to the T2T mPanTro3 chimpanzee reference, and 11,488,040 SNVs from long- (n=5) and short-read (n=13) bonobos mapped to the T2T mPanPan1 bonobo reference (**Fig. 1E-F**, **EDFig. 2**). Next, we calculated SNV heterozygosity from callable sites across individuals. We recovered the expected levels of genetic diversity among chimpanzee populations, with Central and Eastern chimpanzees exhibiting ∼2-fold the SNV diversity of humans in contrast to bonobos and western chimpanzees, which exhibit similar diversity to non-African humans^7^ (**Fig. 1E, EDFig. 2A)**. However, we found that long-read individuals exhibit significantly higher levels of diversity overall (∼6–37% higher on average across populations, **EDFig. 2B**). To explore this further, we compared diversity estimates (π) from long versus short-reads across non-overlapping 10kb genome-wide windows in chimpanzees (**EDFig. 2C-E**). We found that while sequencing methodology does not influence diversity estimates across most of the callable genome, long-read sequenced individuals exhibit hundreds of outlier loci with ∼2-6 fold increased diversity (**EDFig. 2C-E**). These SNV diversity ‘hotspots’ in long-read sequenced individuals are enriched for complex sequences such as segmental duplications, centromeric satellite sequences and subtelomeric regions, where short-reads have reduced mapping fidelity (**EDFig. 2D**). Importantly, such regions are known to exhibit increased mutation rates^32,33^ and play critical biological functions. Principal Component Analysis (PCA) of variants called on either individual long-read haplotypes or jointly with long- and short-read diploid individuals recovered the expected clustering of chimpanzee subspecies (**Fig. 1F, EDFig. 2F**). Together these results highlight that our long-read, physically phased sequencing resources better resolve rapidly evolving regions of the genome still poorly captured with short-reads, and encompass the extensive breadth of chimpanzee and bonobo genetic diversity and structure.

**Figure 2.**
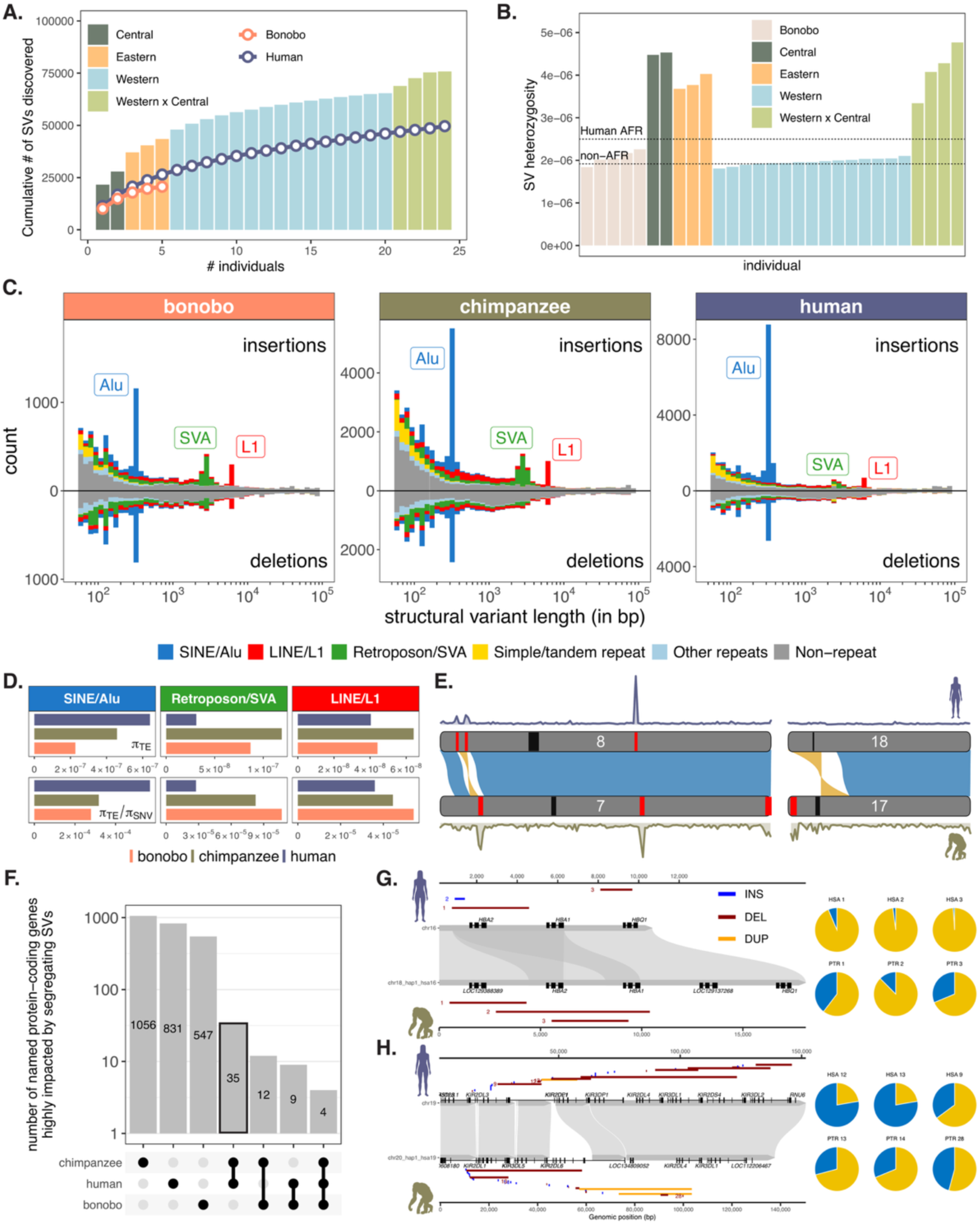
SV mutational patterns and shared SV diversity hotspots across chimpanzees, bonobos, and humans. **(A)** Saturation graph showing the number of structure variants (SVs) discovered with each additional sample. Chimpanzee individuals are represented by bars colored based on their subspecies, whereas humans and bonobos are represented by curves. **(B)** SV heterozygosity in each chimpanzee and bonobo sample, colored based on their species and subspecies. The average SV heterozygosities in human AFR and non-AFR samples from the HPRC-year1 assemblies are shown with dotted lines. **(C)** Length distribution of SVs in bonobos, chimpanzees, and humans. Insertions are shown in the upper part of the plot and deletions are shown in the lower part. SVs are colored by their repeat type, and the three most active TE families in hominids are labeled. **(D)** Nucleotide diversity (π) of full-length TE insertions/deletions and its ratio against π of SNVs. Bars are colored by species. **(E)** Synteny plot showing homologous blocks and large structural rearrangement between human, bonobo, and chimp reference genomes in select chromosomes. Synteny between genomes is shown with blue ribbons when in the direct orientation, and yellow when in the reverse orientation. Centromere locations are represented by black bars. SV hotspots in each species are indicated by red bars. **(F)** Upset plot showing the number of unique and shared protein coding genes highly impacted by SVs across bonobos, chimpanzees, and humans. **(G-H)**. Sequence alignment, SV locations, and allele frequencies at the HBA locus **(G)** and the KIR locus **(H)** in humans and chimpanzees. Grey shades in the middle represent the alignment of human and chimpanzee T2T reference genomes. Gene structures are shown in black boxes at their respective reference. Colored lines represent the location of SVs that segregate in each species (humans on top and chimpanzees on bottom), with colors corresponding to different SV types. Three SVs with the highest allele frequencies are labeled per species per locus, and their allele frequencies are shown by the pie chart, where blue represents the alternate allele and gold represents the reference allele.

### Distinct interspecific SV mutational patterns and shared SV hotspots across chimpanzees, bonobos, and humans

Long-read sequencing provides an unprecedented opportunity to assess structural diversity within and across species. Critically, population-scale long-read datasets allow us to assess segregating structural variants. Here, we employed an ensemble of SV callers to identify and characterize structural variants in chimpanzee and bonobo genomes alongside publicly available human genomes (see methods). By using read-based and assembly-based SV calling approaches with respect to T2T references, we generated and curated species-specific SV catalogues (Sniffles2^34^ and SVIM-asm^35^ for SVs <100kb, and Syri^36^ for SVs >100kb; Supplementary Note 1). Tandem repeats were removed from these SV callsets and analyzed separately^37^. We identified a total of 92,048 SVs across chimpanzees and 23,747 across bonobos respectively (75,799 and 20,628 <100kb), including 54 large inversions (**EDFig. 3**). As a comparative baseline we applied the same pipelines to 47 diverse humans sequenced by the HPRC, and identified a total of 62,915 SVs <100kb, which is comparable to the number of SVs identified by Liao et al^4^.

**Figure 3.**
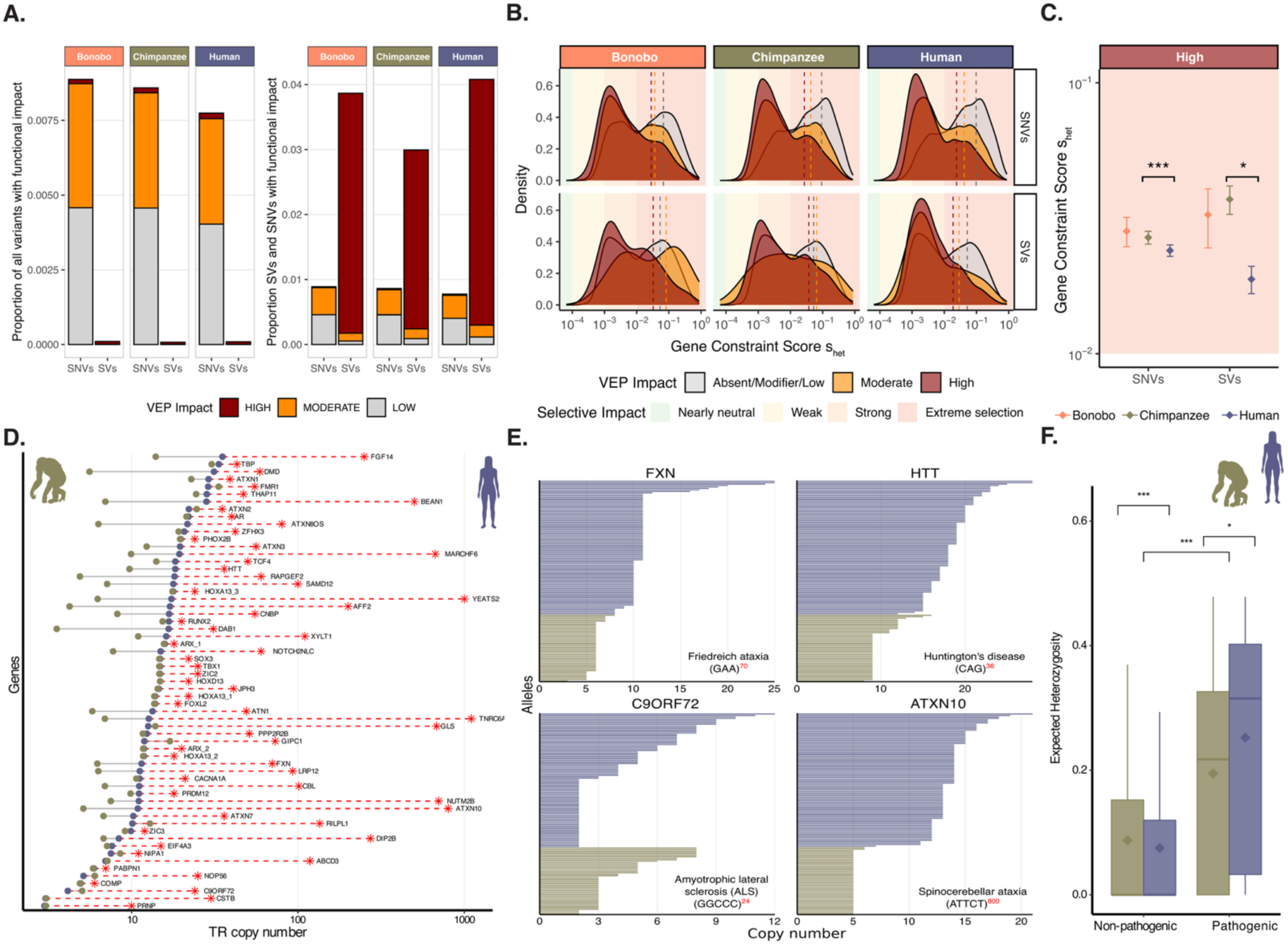
Functional effects, gene constraint and pathogenicity of SVs, SNVs, and TRs across species. **(A**) Stacked-bar proportion of SNVs and SVs in each VEP impact class (HIGH, MODERATE, LOW; MODIFIER excluded) across bonobo, chimpanzee, and human. The left panel shows the fraction of each impact class within all variants (all SNVs + SVs) per species; the y-axis maxes out below 0.01, showing that >99% of all variants are classified as MODIFIER with no predicted functional consequence and that the functional fraction of the genome is dominated by LOW- and MODERATE-impact SNVs. The right panel shows the fraction of each impact class within functional variants only (LOW + MODERATE + HIGH), shown separately for SNVs and SVs. SVs are ∼170–260-fold enriched for HIGH-impact effects relative to SNVs (Bonobo≈260×, Chimpanzee ≈168×, Human ≈203×; see Methods). **(B)** Density distributions of gene constraint (s_het_, log₁₀) for protein-coding genes overlapped by SVs and SNVs of each VEP impact class across species. Dashed vertical lines mark the arithmetic mean s_het_ of each distribution, coloured by VEP class. Background shading indicates selection regimes. In every species and for both variant types, the distribution of genes carrying high-impact variants is visibly shifted toward lower s_het_ (weaker constraint) compared with absent/modifier/low impact variants, and the magnitude of this shift is larger for SVs than for SNVs — confirming that high-impact variants preferentially occur in genes under neutral or weak constraint. **(C)** Mean s_het_ (± SE) of genes carrying HIGH-impact variants in human, chimpanzee, and bonobo, separated by SNVs vs SVs. Asterisks denote significance from two-sided Wilcoxon rank-sum tests vs. human (*** P < 0.001; * P < 0.05). Background shading denotes selection regimes. Chimpanzees carry HIGH-impact SNVs (P = 5.6 × 10⁻⁴) and HIGH-impact SVs (P = 0.044) in genes of significantly higher constraint than humans; the corresponding bonobo–human contrasts are not significant (SNVs P = 0.29, SVs P = 0.17). The effect is magnified for SVs relative to SNVs. **(D)** Mean TR copy number in humans and chimpanzees for TRs located in repeat expansion disorder (RED) genes. Red asterisks represent the pathogenic copy number reported for humans in the Stripy database. **(E)** TR copy number distribution in humans and chimpanzees for four selected RED genes. The associated disorder, TR motif sequence, and pathogenic copy number are indicated for each gene.**(F)** Boxplot of expected heterozygosity for pathogenic versus nonpathogenic TRs in humans and chimpanzees. Asterisks denote significant differences given by the Wilcoxon rank-sum test (*p < 0.05, **p < 0.01, ***p < 0.001).

To further investigate patterns of SV diversity, we compared the cumulative number of SVs <100kb discovered with each additional sequenced sample across species (**Fig. 2A-B**). We found that the saturation curve was highest for chimpanzees, followed by humans, and then bonobos **(Fig. 2A)**. Consistent with SNV patterns, SV heterozygosity was higher in chimpanzees than in bonobos or humans **(Fig. 1E-F**, **Fig. 2B)**. Next, we compared the spectrum of SV sizes and the relative contributions of transposable elements (TEs) to SVs across species (**Fig. 2C**). These distributions were highly similar across species, with three distinct peaks corresponding to the most active TE families in hominids: SINE/Alu, LINE/L1, and SVA. However, we observed marked differences in the relative contributions of these TEs between species; humans exhibited a more pronounced Alu peak while chimpanzees and bonobos exhibited more pronounced L1 and SVA peaks. Increased Alu activity in the human lineage has been previously reported^11^. To more precisely quantify these differences in TE activity we calculated the Tajima’s estimator of *θ*, π, of each of these TE classes across species (**Fig. 2D**). Given that the SNV mutation rate is highly similar between humans and chimpanzees^38^, we can take the ratio of π_TE_ to π_SNV_ to normalize for demographic differences between species and estimate a relative TE mutation rate. These ratios demonstrate that while the human Alu mutation rate is ∼2-fold the chimpanzee and bonobo mutation rates, the SVA mutation rate of chimpanzees and bonobos is ∼3-fold the human rate. L1 activity is similarly higher by about 30% in chimpanzees and 25% in bonobos compared to humans (**Fig. 2D**). Thus, in addition to differences in the diversity of SVs across humans, chimpanzees, and bonobos, which are predominantly driven by demographic processes, these species exhibit distinct TE mutational patterns shaping genetic diversity.

Structural variants are known to form in highly repetitive and unstable regions of the genome including segmental duplications, satellite sequences, and rDNA repeats^32,33,39,40^. Notably, several of these features are shared across recently diverged primate species such as humans and chimpanzees and are thus potentially expected to result in shared “hotspots” of structural variation. To test for such interspecific SV hotspots we calculated the density of SVs across orthologous regions of the genome and found that, indeed, SV density was significantly correlated between humans and chimpanzees/bonobos (**EDFig. 4**, r^2^=0.38 and 0.26 respectively, p<10^−16^). Shared hotspots resolved to several well-known regions of structural complexity including the β defensin locus at 8p23.1 (**Fig. 2E**), the acrocentric short arms, and pericentromeric loci. Nevertheless, as expected given their shorter genetic distance, chimpanzee and bonobo SV density was even more correlated (r^2^=0.47). Chimpanzee- bonobo-specific peaks corresponded to unique features of these primates’ genomes. For instance, the short arm of PTR chromosome 17, (HSA 18) is a chimpanzee/bonobo SV hotspot (**Fig. 2E**, **EDFig. 4A-B**). This locus is the site of one the major cytological differences between humans and chimpanzees, the chr18 pericentric inversion^41,42^ (**Fig. 2E**). Together we identified 109 hotspot loci (34 in chimpanzees, 39 in bonobos, 36 in humans, **EDFig. 4A-B**).

**Figure 4.**
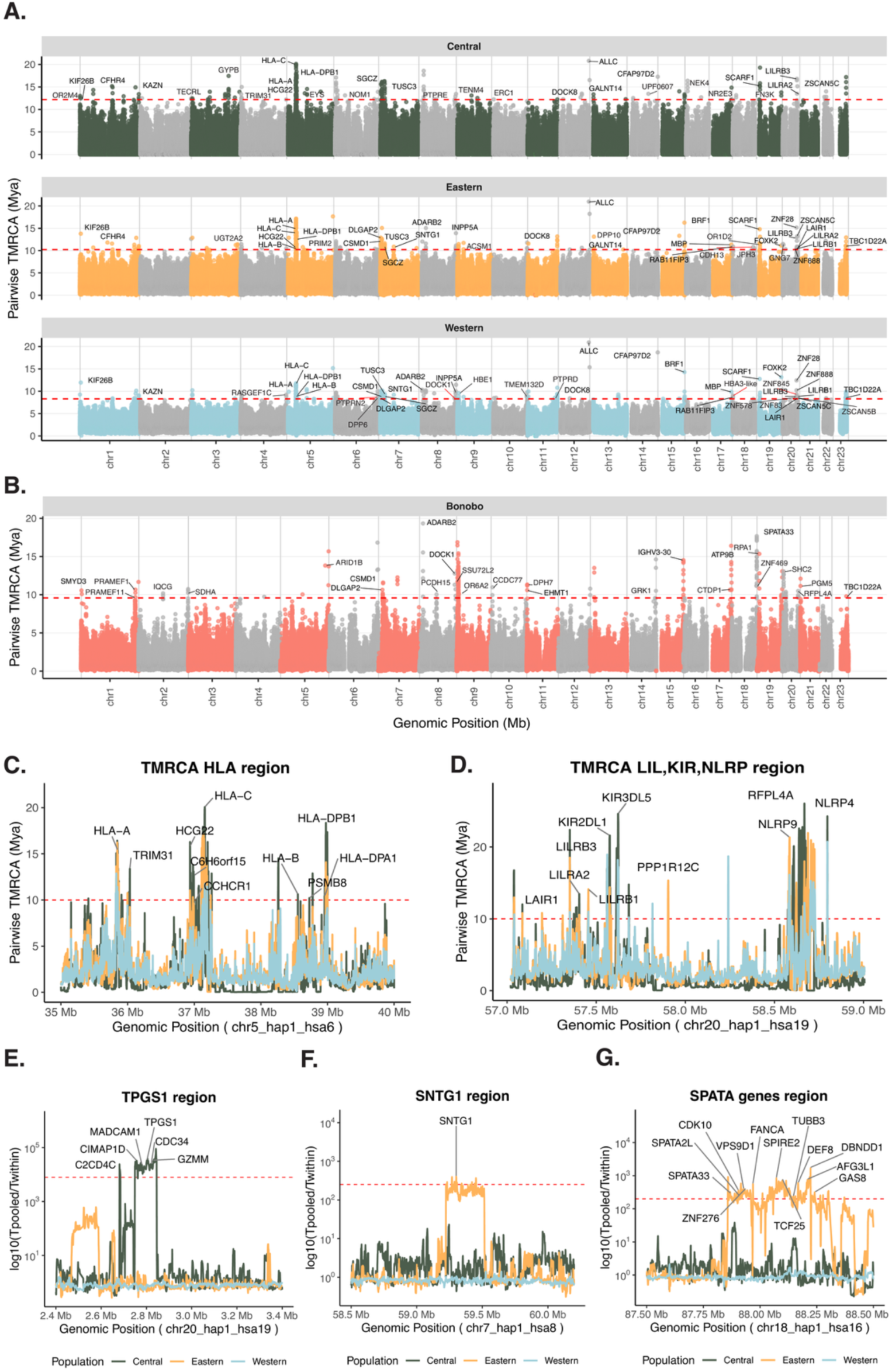
ARG-based detection of balancing and positive selection in chimpanzees and bonobos. (A-B) Manhattan plots of average pairwise time to the most recent common ancestor (TMRCA) in three chimpanzee populations **(A)** and bonobos **(B)** in 1kb windows. Red dotted lines indicate the genome-wide 99.99th percentile. Named genes intersecting with outlier windows are annotated. **(C-G)** Highlighted examples of regions with elevated average pairwise TMRCA **(C-D)** or elevated pooled vs. within-population pairwise TMRCA ratios (*Tpooled/Twithin*) **(E-G)** in chimpanzees. Red dotted lines indicate the genome-wide 99.99th percentile pooled across all populations. Colors correspond to different chimpanzee populations (green: central, orange: eastern, blue: western).

Several SV hotspots intersected genes. We thus sought to characterize the functional impacts of SVs across species, identifying hundreds of lineage specific structural polymorphisms that result in predicted high impact changes to genes (**Fig. 2F**). We also identified dozens of cases in which the same genes exhibited high impact SVs and SNVs across species and chimpanzee populations (**Fig. 2F, Supplementary Table S5-S7**). These include several genes highly impacted by recurrent SVs at the *APOBEC*, *C4A*, *LILRLA/B*, *HBA1/HBA2*, and *KIR* loci (**Fig. 2F-H, Table S5, Supplementary Fig. S1**). In humans, deletions of *HBA1/HBA2* alleles are the primary cause of alpha thalassemias; however, these variants have also been linked to protection against malaria^43,44^, illustrating the complex evolutionary trade-offs shaping genetic diversity. Here, we identify human haplotypes carrying *HBA2* (alpha-2 globin) and *HBQ1* (theta-1 globin) deletions. In contrast, chimpanzees exhibit four distinct haplotypes, each containing 2-4 alpha globin genes (**Fig. 2G**). *KIR* genes, which modulate natural killer cells, have also been associated with malaria susceptibility^45^. We find that the extensive diversity of KIR gene deletions and duplications in humans is mirrored in chimpanzees (**Fig. 2H**), suggesting that similar selective pressures likely driven by host-pathogen coevolution shaped these loci across the *Pan-Homo* lineage. Together, the SV landscapes of humans and chimpanzees exhibit extensive functional convergence as well as unique lineage-specific mutational properties.

### The distribution of segregating functional genetic variation across species

Complete, haplotype-resolved human genome assemblies are increasingly being employed in medical genetics applications given their power to query the full spectrum of genetic variation^46–48^. Such analyses rely on accurate quantifications of the distribution of nonpathogenic variation which can be further contextualized with nonhuman primate genomes. Indeed, recent work has leveraged primate population genetic diversity to develop variant effect prediction models based on the assumption that segregating protein-modifying population-genetic variants in primates are likely to be benign in humans^49^. We thus sought to quantify the impact of SNV and SV diversity in chimpanzees and bonobos in comparison to humans using variant effect prediction (VEP) (**Fig. 3A-C**). While the number of biallelic SNVs alone dwarfs SVs by ∼380-450-fold, SVs were 170-260-fold more likely to exhibit high impact effects across species (**Fig. 3A**), highlighting the functional importance of SVs. These trends are also reflected in the site frequency spectrum where SVs are more enriched at low frequencies compared to SNVs, consistent with stronger purifying selection (**EDFig. 5A**).

**Figure 5.**
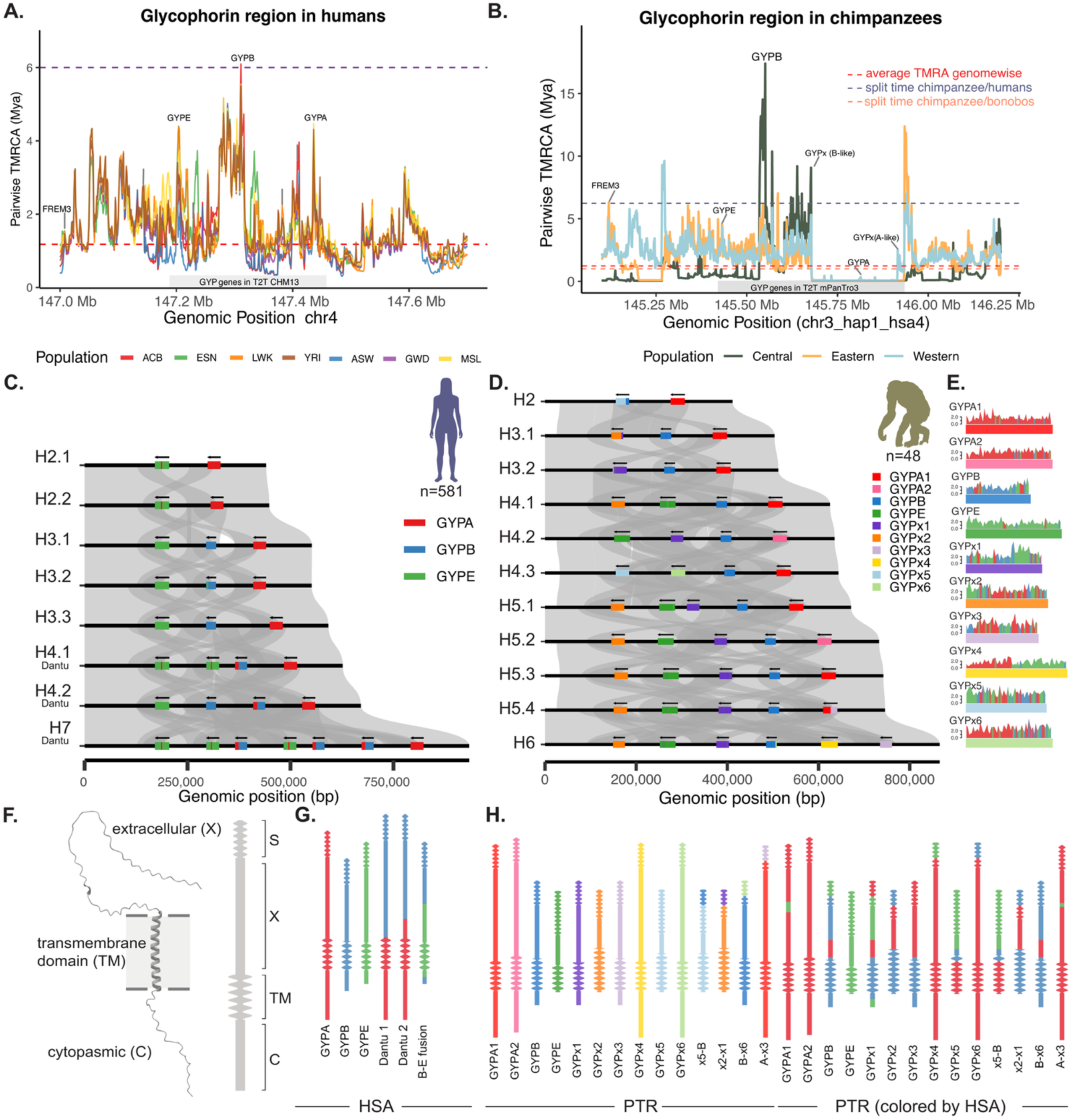
Structural variation and ancient balancing selection at the glycophorin locus in humans and chimpanzees. **(A-B)** Average Pairwise TMRCA estimates across the glycophorin locus in humans **(A)** and chimpanzees **(B)** from long-read sequencing-based phased haplotypes. Dashed lines indicate the average TMRCA genome-wide (red), the human chimpanzee split time (purple), and the chimpanzee bonobo split time (B only, orange). **(C-D)** Stacked pairwise alignments of unique human **(C)** and chimpanzee (**D**) glycophorin haplotypes. Grey ribbons connecting pairs of haplotypes indicate homology relationships. Genes are colored based on homology to “reference” gene annotations thus highlighting both gene fusions and gene conversion patches. Novel chimpanzee genes highly diverged from the reference copies are given unique colors and are named as GYPx1-6. Haplotypes are named sequentially based on the total number of *GYP* genes annotated. Haplotypes including Dantu-like *GYPB-A* gene fusions are labelled. **(E)** Chimpanzee genes are indicated with rectangles with their divergence to their closest human ortholog indicated above, colored by the identity of that ortholog. **(F)** The structure of glycophorin A (excluding the signal peptide) predicted from AlphaFold (left) and a cartoon (right) indicating the cytoplasmic, transmembrane, and extracellular domains as well as the signal peptide. **(G-H)** The structure of the human **(G)** and chimpanzee **(H)** glycophorin proteins with protein domains as described in **(F)** and colored as in (**C-E**).

We next assessed this putatively functional genetic variation in the context of predicted genomic constraint as quantified by s_het_, which estimates the relative fitness reduction of heterozygous loss of function carriers of a gene^50^. As expected, high impact SNVs and SVs were more likely to occur in genes under neutral or weak constraint (**Fig. 3B, EDFig. 5B**). However, the s_het_ of high impact chimpanzee and bonobo variants was significantly higher than high impact human variants (Wilcoxon SNV P = 0.0005, SV P = 0.043; **Fig. 3C**). This effect was magnified in SVs compared to SNVs. Thus chimpanzees and bonobos may carry genetic variation that could be deleterious in humans. We next set out to identify protein-coding genes in chimpanzees and humans that share high impact SVs or SNVs which are also under strong or extreme purifying selection (i.e. s_het_ outliers; **Supplementary Table S8-S9**). We find that highly constrained genes impacted by SVs are significantly enriched for immunity and pathogen response (**EDFig. 5D-F**), including malaria- and HIV related genes (e.g. *BRD9, C4A, MUC19, HBA2, KIR2DL1*). By contrast, while we find several highly constrained genes impacted by SNVs in chimpanzees and humans involved in malaria- and immunity (e.g. *GYPB*, *KIR2DL4*), their functional enrichment is not significant. These results highlight the medical importance and impact of SVs.

One of the most abundant and mutable classes of genetic variation in genomes is found in tandem repeats (TRs) which include short tandem repeats (STRs) with motifs ranging from 1-6bp and variable-number tandem repeats (VNTRs) with motifs >7bp (see companion manuscript Adam et al 2026). TR expansions have been linked to more than 50 human disorders which often manifest as severe central nervous system pathologies^51^. Due to challenges in assaying TRs with short reads, many aspects of their evolution and diversity remain unresolved. We genotyped 61 known pathogenic TR loci in humans and chimpanzees which when expanded in humans result in severe disease^52^ (**Fig. 3D**). The average human TR copy number at these loci was significantly less than the reported pathogenic length in all cases (Wilcoxon P=3.6e-12), as expected given the known deleterious impact of expansions. However, strikingly the average length of these TR loci in chimpanzees was significantly lower than the human average (Wilcoxon P=9.997e-5, e.g. **Fig. 3D-E**). Increased repeat-lengths that are still below the pathogenic length have increased mutation rates and propensity to expand further into pathogenic lengths. This phenomenon, referred to as “anticipation,” results in increased frequency of disease alleles in human populations with longer average nonpathogenic TR lengths^53^. We find that while overall genome-wide TR heterozygosity is higher in chimpanzees than in humans, as expected given their increased genetic diversity, pathogenic TRs have higher heterozygosity in humans, likely as a result of their increased average lengths (**Fig. 3F**). The distribution and dynamics of TR repeat polymorphism in these data are explored in depth in Adam et al^37^. Together our results suggest that human genomes are thus uniquely sensitized to these repeat expansion disorders due to the increased baseline length of these repeats.

### Long-term balancing selection and local adaptation in chimpanzees

A long-standing question in population genetics is how human and great ape genetic variation has jointly and uniquely been shaped by natural selection^7,54–57^. Recent developments in ancestral recombination graph (ARG) reconstruction methods have enabled the discovery and confirmation of long-lived balanced polymorphisms and selective sweeps in humans^58,59^. These approaches infer coalescence trees across the genome thus providing the complete genealogical history of individual loci across a sample of genomes. However, the lack of accurately phased population-scale data has hampered the application of these approaches beyond humans. Here, we leveraged ARGs generated from our 58 physically phased haplotypes using SINGER^58^ to revisit and explore evidence of selection in chimpanzees and bonobos (**Fig. 4, EDFig. 6-7**). Briefly, we computed average pairwise time to the most recent common ancestor (TMRCA) in 1kb windows across populations (**Supplementary Fig. S4**), and focused on regions exceeding speciation times and above the 99.99% percentile of the genome-wide empirical distribution across species (**Fig 4**). We further extended this approach to 218 diverse human haplotypes sequenced by the HPRC-year1 and HGSVC consortia (**EDFig. 6A, Supplementary Table S10**-**S32**).

To identify signatures of long-term balancing selection we specifically looked for deeply coalesced windows in each species harboring variants that plausibly pre-date human and *Pan spp.* divergence (>6 Mya), and thus represent potential transpecies polymorphisms (TSPs). Focusing on the pool of all chimpanzee, bonobo, and human individuals identified several windows harboring genes with deep coalescence predating *Homo-Pan* divergence (14 HSA+PTR+PPA, 6 PTR+HSA, 5 PPA+HSA, 25 PTR+PPA, **Supplementary TableS10-S11**). Refining these analyses to the population-level identified additional loci encompassing a total of 34 deeply coalesced genes in humans and chimpanzees (**Tables S12-S13**). Windows with coalescence times older than the human-chimpanzee divergence were significantly enriched for genes associated with immune-response and host-pathogen interaction (**Supplementary Fig. S5**), spanning several well-established targets of balancing selection such as the *HLA*, *LILR,* and *KIR* genes (**Fig 4A-D, EDFig 6B-E, Supplementary Table S12-S15**)^9,57,60^. We also found several other genes exhibiting exceptionally ancient pairwise coalescence times. These include regions harboring long-lived polymorphisms across humans, chimpanzees, and bonobos (e.g. *ADARB2*, *LILRB3*, *DOCK1*, *SMYD3*, *ADAMT*, *DMBT1*, *TRIM5*, *BTNL2*, *IGFBP7*, *HBE1,* and *MUC5B* among others).

Average Pairwise TMRCA estimates for the HLA region (*HLA-A; HLA-B; HLA-C; HLA-DPB1*) ranged from 10 to 20 Mya in diverse chimpanzees and humans (**Fig 4C, EDFig 5B, Supplementary TableS11-S17**), consistent with previous estimates and evidence of TSPs^9,58^. We note, however, that though we performed strict filtering for regions of the genome susceptible to misalignment and misassembly in our analysis (see Methods), these are the very same loci that are prone to undergo structural rearrangements and are often the targets of the selection and adaptation (**Fig 2**). Loosening these filters identified additional deeply coalesced loci that are shared across species such as the *KIR2DL1* and *NLRP* loci (**Fig 4D, EDFig 6C**). Together these results not only reinforce that balancing selection in *Homo* and *Pan* lineages has repeatedly targeted similar genes involved with immunological pathways, but further show that long-lived balanced polymorphisms are often flanked by or embedded within rapidly evolving structurally complex loci.

To identify signatures of population-specific selective sweeps we next computed the average pairwise coalescence time in the combined sample of haplotypes (T_pooled_) compared to that within each population (T_within_) and focused on the top 99.99th percentile windows overlapped by genes (T_pooled_/T_within_, **Fig 4E-G, EDFig. 7**). In humans, we recovered selection candidates previously identified by Deng et al.^58^ (e.g. *MITF* in AFR), and identified additional candidates such as *KANSL1,* a gene located in the fitness relevant and structurally complex human chromosomal inversion 17q21.31^1–5^ (**EDFig. 7A, TableS33**). In chimpanzees, we found haploblocks exhibiting signatures of selective sweeps overlapping several genes involved with spermatogenesis, brain development, and infectious disease (**Fig 4E-G, EDFig. 7B, Supplementary Table S34**). Central chimpanzees, in particular, exhibit signatures of a recent selective sweep in a haploblock harboring *TPGS1* (**Fig 4E**), a biomarker for leprosy in humans^61^. Notably, clinical manifestations of naturally acquired leprosy have been recently described in western chimpanzee wild populations in Guinea-Bissau and Cote d’Ivoire^23^, but its prevalence in Central populations has not yet been documented. In Eastern chimpanzees, top candidates included haploblocks harboring several genes in the sub-telomeric region of chromosomes 18 and 20 such as *SPATA33,* which is implicated in sperm motility. We also identified a region harboring *SNTG1*, a gene involved in brain development that underwent a recent selective sweep in humans^62^ and harbors anthropoid-specific constrained regulatory sequences^63^. In Western Chimpanzees, top candidates included *HLA*-related genes alongside olfactory receptors and genes implicated in neurodevelopment (**EDFig. 7B, Supplementary Table S34)**. Together, these results highlight the utility of haplotype-resolved long-read genome assemblies for ARG-based population genetic inference in nonhuman species.

### Ancient balancing selection and recurrent structural variation at the malaria-associated glycophorin locus in humans and chimpanzees

We identified several loci exhibiting extensive structural variation in humans and chimpanzees overlapping genes implicated in malaria resistance, including the glycophorin locus (**EDFig. 5D-F, Supplementary Table S5-S7 and Fig. S5)**. Glycophorin genes, including *GYPA*, *GYPB* and *GYPE,* encode for highly glycosylated erythrocyte transmembrane proteins which determine the MNS blood group antigens^64^. While glycophorins interact with several different pathogens they play a critical role as receptors enabling erythrocyte invasion by *Plasmodium falciparum,* the major cause of malaria in Africa^65,66^. *GYPB* and *GYPE* originate from a duplication of *GYPA* at a single ∼300kb locus in the ancestor of African great apes (gorillas, chimpanzees, bonobos, and humans). Short read sequencing data, FISH, and array typing^65,67^ have all demonstrated that the locus exhibits extensive structural polymorphism in humans. Furthermore, the Dantu blood group antigen, which is protective against severe malaria and largely found in East Africa, is the result of a distinctive structural variant resulting in a fusion between the *GYPA* and *GYPB* genes^65,68^. More recently, variants at this locus have been associated with local adaptation in wild chimpanzee populations^55^. However, despite long standing interest into this medically important region, the complete structure and sequence of the glycophorin locus remains unresolved outside of reference genomes.

To better understand the evolution of the glycophorin genes we first estimated average pairwise TMRCA in 1kb windows across the locus in chimpanzees and 581 diverse, physically phased human haplotypes (see methods, **Fig. 5A, B, EDFig. 8A-D**). In humans GYP genes exhibit pairwise coalescence times ranging from ∼4-6mya, significantly higher than the estimated genomes wide ∼1 Mya coalescence time (**Supplementary Fig. S4**). Chimpanzee TMRCAs are even more extreme with several peaks exceeding the *Homo-Pan* divergence supporting previous work highlighting the likely impact of long term balancing selection at this locus^57^. We next sought to determine the structural diversity of the glycophorin locus in both species (see methods). Unique haplotype structures were identified by first constructing a pangenome variation graph and then clustering haplotypes based on their pairwise basepair-level Jaccard similarity. To further dissect differences between haplotypes we annotated genes across each haplotype and assigned each 500bp tiled segment of a gene an identity based on its closest match to reference genome annotations. Unique haplotypes thus represent both novel gene configurations, as well as unique cases of gene conversion and gene fusion events.

Across 581 human haplotypes we identified 8 unique human glycophorin structural configurations each spanning 2-7 *GYP* copies (**Fig. 5C**). We find that ∼97% of human haplotypes carry the reference H3.1 haplotype, with the remaining structural diversity at GYP largely concentrated in African and African-admixed populations (**EDFig 7E**). We identified three haplotypes exhibiting *GYPB-A* gene fusions occurring at low frequency resembling those reported in malaria-protective Dantu gene fusions. We refer to these haplotypes, each containing 4-7 total *GYP* genes as H4.1_Dantu1_, H4.2_Dantu2_ and H7_Dantu1_. The longest of these, H7_Dantu1_, contains 3 *GYPA-B* fusion genes, 3 *GYPE* genes, and a single *GYPA.* The H4.1_Dantu1_ haplotype differs from H7_Dantu1_ only in the loss of 2 intervening *GYPA-B* genes and a *GYPE* gene in a manner consistent with non-allelic homologous recombination (NAHR). Thus we hypothesize H7_Dantu1_ and H4.1_Dantu1_ share the same origin. However, the H4.2_Dantu2_ haplotype exhibits an independent gene configuration similar to the reference haplotype with the addition of a *GYPA-B* fusion gene between *GYPB* and *GYPA* genes. Remarkably, while in H4.1_Dantu1_ and H7_Dantu1_ the *A* portion of the *A-B* fusion is ∼8.5kb, in the H4.2_Dantu2_ haplotype the *A* portion of the *A-B* fusion is ∼10kb long. We conclude that this fusion is thus likely an independent event. Notably, while the H7_Dantu1_ haplotype is found in an individual with African ancestry, the H4.1_Dantu1_ haplotype is identified in an individual from the United Arab Emirates (UAE) and the H4.2_Dantu2_ haplotype is found in a a Peruvian individual with no evidence of African admixture over this locus (**EDFig. 8E**). Alongside the Dantu *A-B* fusions we also identified an *E-B* gene fusion in the H3.2 haplotype in a single UAE individual, similar to a “hybrid gene” associated with a protease resistant antigen recently reported at low frequency in Japanese populations^69^.

Across our 48 chimpanzee haplotypes we identified 11 distinct glycophorin haplotypes each containing 2-6 *GYP* copies, a ∼17-fold increase in haplotype diversity compared to humans (**Fig. 5D**). Chimpanzee *GYP* genes were also substantially more diverse than human genes as well. We identified 10 different *GYP* genes including single copies of *GYPB*, and *GYPE* and 2 distinct, highly diverged *GYPA* genes (∼2.4% divergence). However, alongside these canonical *GYP* genes we also identified 6 additional novel *GYP* genes (designated *GYPx1-6*). To better understand the origin of these genes we assigned each 500bp tiled segment of these genes to its closest matching human *GYP* paralog (**Fig. 5E**). These tiled homology assignments revealed that each *GYPx* gene is made up of a patchwork of *GYPA/B/E* segments, likely the result of ectopic recombination and interlocus gene conversion over millions of years. Some of these chimeric architectures resemble ancient gene fusion events, such as *GYPx4,* which is an ancient fusion between *GYPA* and *GYPE.* In addition to these ancient gene conversion and fusion events, we also identified 5 more recent gene fusion events in chimpanzees, resembling those identified in humans. Thus, chimpanzees and humans exhibit parallel evolutionary histories at the GYP locus characterized by extensive gene duplication, gene conversion, and gene fusion events which have shuffled the sequences of these genes. However, glycophorin genetic diversity in chimpanzees is much older, potentially reflecting more ancient selective pressures as a result of the prolonged contact of chimpanzees with malaria vectors.

A recent study of wild-derived chimpanzee exomes discovered variants associated with adaptation to forest cover intersecting the glycophorin locus and other malaria-associated loci^55^. This work concluded that forest cover-associated alleles resulted in stop codon gain, loss of function variants in *GYPA*. Our complete haplotype assemblies enabled us to revisit this result and determine that the putative stop-codon gain is a likely short-read mapping artifact (see **Supplementary Note 2 and Supplementary Figure S6**). Instead, we find forest-cover associated variants are likely associated with the H6 chimpanzee haplotype (**EDFig. 8F**). This haplotype includes several recent and ancient GYP fusions including 4 of the newly described *GYPx* genes. We used haplotype deconvolution^3,70^ to genotype the complex glycophorin haplotypes present in 96 short-read sequenced chimpanzees and bonobos (**EDFig. 8F**). These results reveal that glycophorin haplotypes exhibit strong population stratification and geographic differentiation among chimpanzee subspecies. The H6 haplotype, in particular, was found only in Western chimpanzees (18% frequency). Together, these results show that haplotype-resolved assemblies enable novel insights into local adaptation in wild chimpanzee populations and further the utility of precious chimpanzee genomic resources.

Glycophorins are single transmembrane domain proteins with disordered cytoplasmic and extracellular domains and a signal peptide sequence which is cleaved (**Fig. 5F**). We annotated these structural features onto human and chimpanzee *GYP* coding sequences to understand the functional impact of gene fusion and gene conversion events (**Fig. 5F, H**). As has been previously reported, the Dantu1 protein (found in H4.1_Dantu1_ and H7_Dantu1_) consists of *GYPA*-derived cytoplasmic and transmembrane sequences fused to a *GYPB*-derived extracellular domain (**Fig. 5G**). The newly discovered Dantu2 protein however maintains 13 additional amino acids of *GYPA* extracellular sequence. The GYPE-B fusion protein exhibits a GYPE-derived transmembrane domain with partial contributions to both the extracellular and cytoplasmic domains from *GYPB* (**Fig. 5G**). Recent chimpanzee gene fusion events mostly impact the signal peptide sequences. However, the ancient gene fusion and conversion events have extensively shuffled protein sequences with respect to the canonical GYP proteins including multiple GYPA-E and GYPB-A fusions (**Fig 5H**). Together these results highlight the importance of structural variation across taxa in generating functional novelty through gene duplication, fusion, and conversion.

## Discussion

Here we sequence and assemble chimpanzee and bonobo genomes from four of the five recognized subspecies and species of the *Pan* genus. This resource captures unprecedented sequence and structural diversity providing access to previously intractable genomic regions, including rapidly evolving hotspots of structural variation and adaptation. Inclusion of these rapidly evolving loci increases estimates of genome-wide heterozygosity by as much as 37% compared to short read sequencing in some populations, highlighting the extensive genetic diversity missed by short-reads. Critically, these high-diversity regions are biologically and evolutionarily important, often having been the targets of selection for millions of years. Population-scale comparative long read sequencing enables some of the first comprehensive analyses of these loci.

Long-read sequencing of chimpanzee and bonobo genomes also enabled us to perform comparative population-scale analyses of biomedically relevant regions such as tandem repeat expansion disorder loci. We find that these disease-associated tandem repeat loci are systematically longer, and closer to pathogenic lengths in human populations compared to chimpanzee populations. This highlights how human genomes are uniquely sensitized to predispose us to certain genetic diseases. Similarly, the genome-wide interspersed distribution of segmental duplications predisposes humans and other great apes to micro-deletion/-duplication disorders^71^. Nevertheless, it is important to consider that these repeat expansion disorders were discovered and described in humans^51–53^. Thus, ascertainment of disease-associated TR loci in chimpanzees would likely exhibit the opposite result. This finding highlights that across the tree life species likely exhibit lineage-specific “anticipation” for diseases harbored in their genomes. While the “genetic load,” or the burden of deleterious variation in individuals, is often contrasted between species and populations, these results highlight that the mutation propensity of not-yet-realized deleterious variants is another important factor that can result in fitness differences.

Recent advances in ARG-based inference have enabled ever more powerful insights into demography and selection across the genome, yet rely on phased data^58,59^. Here we leverage diverse physically phased chimpanzees and bonobos to construct genome-wide ARGs and identify putative targets of long-term balancing selection. Many of these coincide with structurally complex regions, including the well-studied glycophorin locus. In humans, the Dantu blood group antigen has been shown to reduce the risk of severe malaria by up to 79%^66^. The genetic basis of this protection is through the generation of a novel fusion gene between *GYPA* and *GYPB*. This hybrid receptor has been shown to increase erythrocyte membrane tension, impairing the ability of *P. falciparum* merozoites to invade the cell^66^. We describe the complete sequence and structure of *GYPA-B* fusion genes and the haplotypes upon which they exist. We find that there are likely at least two independent origins of Dantu fusion genes as well as additional *GYPA-E* fusions that may be of functional relevance. Chimpanzees are also susceptible to malaria^72^ and have coevolved with this parasite for millions of years. We find that, mirroring the evolutionary innovations in humans, chimpanzees exhibit prolific gene duplication and fusion at the glycophorin locus creating extensive novelty. In addition to higher overall haplotype diversity, chimpanzees also harbor several novel GYP genes. We find that these genes are the result of ancient fusion and conversion events resulting in protein structures that resemble chimeras of the human protein diversity. While it is not known if these novel genes confer similar surface tension phenotypes in erythrocytes, they strikingly resemble the same patterns of diversification as humans, albeit over longer timescales, underscoring the importance of gene duplications and fusions to rapidly create evolutionary novelty for adaptation. We further identify additional repeated signatures of selection and recurrent structural polymorphisms across taxa. This is consistent with recent comparisons of high-quality primate reference genomes spanning 50 million years of evolution, which similarly identified extensive recurrent structural variation^73^. These patterns highlight that adaptations across vast evolutionary distances are often fueled by genetic substrate harboring extensive structural complexity.

The remarkable contiguity of the chimpanzee and bonobo genomes presented here enables comparative analyses of many rapidly evolving complex loci. However, regions of extreme complexity such as centromeres, rDNA repeats, and the Y-chromosome remain elusive to population-scale comparative approaches without complete T2T resolution. The limited sample sizes for some clades in our resource also limits our ability to fully characterize natural selection across populations. Signals of population-specific balancing and directional selection in immune response loci (e.g. HLA genes) and genes implicated in neurological development, respectively, may be tied to the selective pressures of disease landscapes, specifically SIV^74^, and habitat-specific socioecologies^18,75^. This resource alone does not fully reflect the extensive ecological diversity and fine-scale geographic structure of locally adapted wild populations, most notably in unsampled Nigeria-Cameroon chimpanzees (*P. t. ellioti*) and other populations known to exhibit unique signatures of local adaptation and resistance to disease across diverse habitats^30,31,55,56,76–78^. Nevertheless, this work demonstrates how nonhuman primate pangenomes can be integrated with existing datasets to shed light into the evolution and diversity of these species, while simultaneously generating novel insight into the evolution of complex genomic architectures in humans. As massive-scale sequencing efforts continue to transform our understanding of human genetic variation, we envision future population-scale long-read sequencing initiatives across the tree of life as critical to contextualize this diversity and understand the processes driving genome evolution.

## Material and Methods

### Datasets

Newly generated long-read sequencing data for chimpanzees and bonobos featured in this work are named PANPAN (*Pan* spp. pangenome project; n = 5 bonobos; n=24 chimpanzees, **Fig. 1A**). Several additional different datasets were assessed in this manuscript from publicly available sources. Human long-read sequencing data includes the previously generated datasets by the HPRC (Human Pangenome Reference Consortium) and HGSVC (Human Genome Structural Variation Consortium)^4,26^; and additional complementary haplotypes from publicly available long-read genomes (https://github.com/lh3/OpenHGL). Chimpanzee short-read sequencing data includes publicly available data from NCBI bioprojects PRJEB15086 and PRJNA189439.

### Chimpanzee and bonobo sample selection and cell culture

All samples were selected from available lymphoblastoid (LCLs) and fibroblastoid cell lines from the Integrated Primate Biomaterials and Information Resource (IPBIR) at the Coriell Institute for Medical Research. We first generated low-coverage Illumina short-read resequencing data for all of these cell lines (https://github.com/sudmantlab/panpan) and used this preliminary data to select a subset of a individuals for long-read PacBio HiFi sequencing, prioritizing samples that maximized population diversity and structure, favoring wild-born individuals, and excluding cell lines exhibiting slow-growth phenotypes. Selected cell lines were then expanded to a total culture size of 3×10⁶ cells. The cell line expansions were derived from the original expansion culture to reduce the number of passages and minimize culturing time. Cells were washed in PBS and flash-frozen as dry cell pellets of 1×10⁶ cells per vial. All data are outlined in **Supplementary Table S1**.

### DNA isolation and sequencing

#### HiFi PacBio Sequencing

We isolated high molecular-weight DNA (HMW DNA) from these samples using Circulomics CBB kit (102-573-600) from a frozen cell pellet (10e6 cells). Elution was performed overnight at room temperature. DNA quantity, purity and integrity were checked at different steps and at the end of the extraction protocol. DNA quantity was checked on a Qubit Fluorometer I with a DNA High Sensitivity (DNA-HS) Qubit assay (Invitrogen), and sizes examined on a Fragment Analyzer or FEMTO pulse (Agilent Technologies) using a Genomic DNA 165kb kit. Purity ratios were assessed with NanoDrop. A total of 54.8 micrograms of DNA (274 ng/uL in 200 uL volume, over 50kb length, and purity ratio 260/280: 1.82, 260/230:2.0) was used as input for library preparation. Samples were sequenced by HiFI in two major batches on the Sequell II and Revio platforms.

#### Sequel II sequencing

A starting amount of 4-5 ug HMW gDNA was sheared to a target size of 20-30 kb using a Megaruptor 3 instrument (Diagenode). The sheared DNA underwent size selection using a Pippin HT instrument (Sage Science) to target a size range of 15-22 kb. Following size selection, the DNA was used for CCS (Circular Consensus Sequencing) library preparation using the SMRTBell Express Template Prep Kit 2.0 and Enzyme Cleanup Kit 1.0 (PacBio). Each library was barcoded using PacBio Barcoded Overhang Adapters. Post-library preparation, the concentration of the DNA stock was measured using the DNA-HS Qubit assay, and the DNA size was estimated using the Fragment Analyzer or Femto Pulse. Sequencing was conducted on a Pacific Biosciences Sequel IIe instrument, using version 2.0 sequencing reagents and operating on control software version 10.1.0.119549, with a movie collection time of 30 hours per 8M SMRT Cell with no pre-extension and with adaptive loading. CCS/HiFi reads were generated from the initial subread data using the CCS program version 6.0.0 within PacBio SMRTLink version 10.1.0.119588. Fastq sequences were extracted from BAM files using SMRTLink 10.1.0.119588. These are labelled as PR* samples.

#### Revio sequencing

A starting amount of 1.5 ug of HMW DNA was used for library preparation following the Pacific Biosciences SMRTbell prep kit 3.0. The Megarupter (Diagenode) was used for shearing and a BluePippin Instrument (Sage Science) was used for size-selection for fragments over 10-50kb. The library was run on a PacBio Revio Instrument using Revio SMRT cells (102-817-900) sequencing reagents. Sequencing was performed with SMRTells running instrument control software version V13 and a movie collection time of 24 and 30 hour movie hours per SMRTCell with a 2-hour pre-extension and adaptive loading enabled with V13. CCS/HiFi reads were generated from the initial subread data using the ccs program version 8 within PacBio SMRTLink version 13.

#### Hi-C sequencing

Hi-C data was additionally generated for some PR* and all AG*-labeled samples. Hi-C libraries were generated from 1M cells at Passage 3 using the Illumina HiC kit (Dovetail genomics); libraries were submitted for quality control and sequencing on the Illumina NovaSeq 6000 platform (Novogene).

#### ONT sequencing

Ultra-long-(UL-)ONT were generated according to a previously published protocol (Logsdon, protocols.io, 2020). Briefly, 3-5 x 107 cells were lysed in a buffer containing 10 mM Tris-Cl (pH 8.0), 0.1 M EDTA (pH 8.0), 0.5% w/v SDS, and 20 mg/mL RNase A (Qiagen, 19101) for 1 hour at 37°C. 200 ug/mL Proteinase K (Qiagen, 19131) was added, and the solution was incubated at 50°C for 2 hours. DNA was purified via two rounds of 25:24:1 phenol-chloroform-isoamyl alcohol extraction followed by ethanol precipitation. Precipitated DNA was solubilized in 10 mM Tris (pH 8.0) containing 0.02% Triton X-100 at 4°C for two days. Libraries were constructed using the Ultra-Long DNA Sequencing Kit (ONT, SQK-ULK001 and ULK114) with modifications to the manufacturer’s protocol. Specifically, ∼40 ug of DNA was mixed with FRA enzyme and FDB buffer as described in the protocol and incubated for 5 minutes at RT, followed by a 5-minute heat-inactivation at 75°C. RAP enzyme was mixed with the DNA solution and incubated at RT for 1 hour before the clean-up step. Clean-up was performed using the Nanobind UL Library Prep Kit (Circulomics, NB-900-601-01) and eluted in 225 uL EB. 75 uL of library was loaded onto R9 and R10 flow cells for sequencing on the PromethION, with two nuclease washes and reloads after 24 and 48 hours of sequencing. Data was basecalled with guppy/6.3.7 using the sup basecalling model.

### Genome assembly

#### Read processing and genome assembly

PacBio HiFi CCS reads were adapter-filtered with HiFiAdapterFilt-v2.00^79^. Paired-end Hi-C reads were processed with Trimmomatic v0.35-6 to remove adapter sequences and low-quality bases (settings *ILLUMINACLIP:<u>TruSeq2-PE.fa:2</u>:40:15:SLIDINGWINDOW:5:20*). For samples with HiFi reads only, we ran hifiasm-v.0.2.2 to generate haplotype-resolved contigs (*.hap1.p_ctg.gfa, *.hap2.p_ctg.gfa) and a primary contig set (*.p_ctg.gfa). For samples with both HiFi and paired end HiC data, hifiasm was run with Hi-C–integrated mode providing both the CCS reads and the trimmed Hi-C reads as input, and purging duplicates using the −l2 option (*.hic.hap1.p_ctg.gfa, *.hic.hap2.p_ctg.gfa, *.hic.p_ctg.gfa). For samples with additional ONT data, haplotype-resolved and diploid assemblies were built with Verkko-2.2.1, combining the adapter-filtered HiFi reads, ONT reads and trimmed Hi-C R1/R2. Assembly graphs were converted to FASTA with gfatools-v0.5.5 (https://github.com/lh3/gfatools).

#### Quality control

For each individual, k-mers were counted from the adapter-filtered HiFi reads with Jellyfish-v2.3.1^80^ and GenomeScope-v2.0^81^ was run on individual fastq files to obtain per-sample estimates of heterozygosity, sequencing coverage, repeat content and genome size. Meryl-v1.3^82^ databases were built per sample (meryl count k=21) and merged across runs (meryl union-sum). The merged database was used by Merqury-v1.3^82^ to estimate consensus quality values (QV) and k-mer completeness for (i) the primary-contig HiFi-only assembly (*.consensus) and (ii) respective hap1 and hap2 assemblies (*.haplotig). Assembly contiguity statistics (NG50, auN/auNG) were computed using both a costume made tool (https://github.com/sudmantlab/assemblystats) and with gt seqstat -contigs -genome <genome_size> (GenomeTools-v.1.6.2; https://github.com/genometools/genometools), where the genome size was taken from GenomeScope2 model fit.

#### Reference scaffolding

All haplotypes and primary assemblies for all species in this study were additionally reference-scaffolded with RagTag-v2.1.0^83,84^ (-C -u --mm2-params ‘-x asm5’ --unimap-params ‘-x asm5 -t 20) against conspecific telomere-to-telomere references: chimpanzee T2T-mPanTro3, bonobo T2T-mPanPan1, and human T2T-CHM13. The - C flag concatenates unplaced contigs into a single ‘Chr0’ scaffold, which was subsequently excluded; remaining scaffolds were stripped of the _RagTag suffix and name-sorted with seqkit-v2.10.0 (https://github.com/shenwei356/seqkit). This allowed converting contigs into the same chromosome naming convention for downstream analysis. Because mPanTro3 and mPanPan1 derive from male individuals with fully resolved X and Y chromosomes, scaffolded assemblies were aligned back to these references with minimap2 (-x asm5 -c --eqx --cs --secondary=no) and the resulting PAF inspected with pafR (https://github.com/dwinter/pafr); coverage patterns confirmed reported sex for Coriell-derived samples and further identified sex-chromosome contigs in each query assembly for unknown individuals (all featured in **Supplementary Table S1**)

#### Genome completeness and T2T status

To further evaluate the assembly quality, we quantified gaps and telomeric repeats for all chromosomes across each haplotype. Gaps, defined as unresolvable genomic regions resulting from complex graph structures or sequencing/library preparation dropouts, were identified using a custom awk script. Telomeres (telomere motif: TTAGGG) were identified using the telo function in seqtk v1.4-r130(https://github.com/lh3/seqtk). Chromosome completeness was assessed based on the presence of flanking telomeres and the absence of internal gaps. Each chromosome was categorised into one of six hierarchical states: (1) T2T (gapless with two telomeres), (2) gapless with one telomere, (3) gapless with zero telomeres, (4) with gaps and two telomeres, (5) with gaps and one telomere, and (6) with gaps and zero telomeres. These metrics were aggregated per haplotype and compared across species and subspecies/populations. The distribution of gaps, telomere counts, and T2T-status chromosomes was benchmarked against established T2T reference assemblies (mPanPan1 and mPanTro3) to validate the continuity of the PANPAN dataset (**Supplementary Table S2-S4**)

### Variant discovery

#### Read and assembly alignments to T2T references

All reference-based alignments used the three telomere-to-telomere assemblies (T2T-CHM13, mPanTro3 and mPanPan1) as targets, with each sample mapped to its species-specific reference. Illumina short-reads were quality-trimmed with Trimmomatic-v0.40 (SLIDINGWINDOW:4:20 MINLEN:30) and aligned with BWA-MEM-v0.7.18^85^; alignments were flag-filtered with samtools-v.1.21^86^ (-q 15 -F 780, removing unmapped, mate-unmapped, secondary, QC-fail and duplicate records), deduplicated with Picard MarkDuplicates-v2.27.2 (https://broadinstitute.github.io/picard/), and indel-realigned with GATK-v3.5 (http://www.broadinstitute.org/gatk/). PacBio HiFi long-reads were aligned with winnowmap-v2.03^87^ (-x map-pb -Y -L --eqx --cs) using high-frequency k-mer mask built with meryl (k=15), filtered at -q 10, tagged with per-run read groups and merged per individual with samtools-v.1.21^86^. Haplotype-resolved assemblies were aligned with minimap-v.2.28-r1209 (-cx asm5 –cs) and the resulting PAFs were sorted by target and position prior to variant calling. Coverage per-bam was assessed with deepTools-v.3.5.5^88^.

#### SNV calling

From the read-based alignments, single-nucleotide variants (SNVs) were jointly called per species with bcftools-v1.21^86^ (bcftools mpileup -q 30 -Q 20 -a AD,DP,SP followed by bcftools call -m -f GQ,GP), merging and sorting long- and short-read–derived bam files into a multi-individual .vcf per species with respect to conspecific T2T reference. All-sites VCFs were also filtered with vcftools-v0.1.16 (--remove-indels --max-alleles 2 --max-missing 0.9 --maxDP 200), with per-site mean-depth bounds set per species to bracket the panel-wide mean coverage (--min-meanDP 24 --max-meanDP 37 across the 83 long- and short-read chimpanzee individuals; and --min-meanDP 10 --max-meanDP 55 for bonobos for 18 long- and short-read bonobos). A biallelic-SNP subset was generated with the same depth/missingness thresholds plus --min-alleles 2 --minQ 30 --minGQ 20. All-sites VCFs generated with long-read and short-read mapped to T2T reference genomes were then used as input to pixy-v2.0.0.beta8 to estimate per-site nucleotide diversity per individual (heterozygosity bp -1). We also ran pixy to estimate per-window nucleotide diversity (π) across 10 kb non-overlapping windows in long-read sequenced individuals compared to short-read sequenced individuals. Genomic features (centromeres, telomeres, centromeric satellites, segmental duplications, tandem repeats, and short-read accessibility mask) were intersected with the window grid, so that each window carried its feature annotation. For each window we computed the π_long / π_short ratio after discarding windows with π = 0 in both sets. Outlier loci were defined as the top 1% of windows by this ratio per population, and per-feature fold changes were computed as the within-population mean π_long / π_short across all windows overlapping each feature class. Biallelic SNP subsets of the joint read-based VCFs (combining long- and short-read–sequenced individuals) were used as input for principal-component analysis with PLINK-v1.9.0-b.7.7^89^ (bcftools view --types snps --min-alleles 2 --max-alleles 2). In parallel, an assembly-based SNV call set was generated for the long-read–sequenced individuals using paftools.js call (minimap2) alignments of each haplotype-resolved assembly to its species-specific T2T reference, and the resulting VCFs were also used as input for PCA. The 2 types of read-based (all-sites and bi-allelic snps .vcf for chimpanzees and bonobos long- and short-reads) and assembly-based (paftools call .vcf for chimpanzees and bonobos long-reads) callsets were then used for downstream analysis.

### SV calling

**SVs<100kb** were called from HiFi reads and from assemblies. First, we used the findTandemRepeats script in pbsv-v2.9.0 https://github.com/PacificBiosciences/pbsv to annotate tandem repeat regions in reference assemblies. We passed these regions to the --tandem-repeats flag in sniffles-v2.0.7^34^ to call SVs from HiFi reads aligned to species-specific references for each sample separately. We then used sniffles to combine individual .snf files into a multiple-sample vcf file for each species. We filtered the vcf for variants that passed built-in filters, that have precise breakpoint locations, and that are between 50bp and 100kb in length. We removed insertions and deletions without an alternate allele sequence or with inconsistent svlen and seqlen values. We removed breakend variants (BND), and filtered out any sites with extremely high read depth (top 5%) to avoid artefacts caused by paralogous alignment (unless the SV is a duplication). Separately, we called SVs from haplotype-resolved assemblies aligned to the reference genome using svim-asm-v1.0.3^35^ under the diploid mode. Resulting individual-level vcf files were merged with bcftools-v1.9 merge -m none followed by truvari-v4.2.0 with the following parameters: collapse --chain -r 1000 --sizemin 50 -p 0.9 -P 0.9. Since missing genotypes in the merged vcf file are likely not truly missing, they were converted to homozygotes for the reference allele using bcftools +missing2ref. Similar to sniffles, we filtered for SVs that passed the built-in filter, that are between 50bp and 100kb in length, and that are not of type BND. To merge the two SV callsets from different callers, we first used bcftools merge - -force-samples followed by travari collapse with the same settings as the merging step across samples. The only exception is that for inversions and duplications, the -p flag which establishes a sequencing similarity threshold for merging was set to 0 instead of 0.9 because sniffles does not report the inversion and duplication sequences in its vcf. For variants that are supported by both callers and those that are only supported by svim-asm, genotypes reported by svim-asm were used for the final vcf file. For variants supported by sniffles only, sniffles’ genotypes were used. Furthermore, we extracted the insertion and deletion sequences from the merged vcf file and ran repeatmasker-v4.1.2-p1 to annotate their repeat type. An SV is considered to be of a certain repeat type if more than 80% of its sequence is annotated by repeatmasker and if the top repeat type occupies more than 80% of the annotated part of the sequence. Otherwise it is labeled as a “non-repeat”. As a final filtering step, any insertions and deletions that are not labeled as a transposable element and intersect with tandem repeat regions of the reference genome are filtered out to reduce the noise at these complex regions. Individual heterozygosity and population-level theta estimates of SVs were computed from the final vcf file using the full genome size as the denominator. Theta estimates of TEs only included full-length TEs (>250bp for Alu, >1800bp for SVA, >5000bp for L1). The density of SVs across the genome was estimated in 1Mb non-overlapping windows, and hotspots are defined as windows with SV density three standard deviations above the mean.

**SVs>100kb** were identified across all samples by initially aligning individual haplotypes to their respective reference genomes using minimap2. SVs were called using SyRI v1.7.0^36^, which identified inversions, duplications, translocations, insertions, deletions, and highly diverged regions (HDR; symmetric low-quality or missing alignments). Overlapping inversions within each species population were identified, and the encompassing genomic regions, including 20 kb of flanking sequence, were extracted from each genome. These regions were independently aligned to their reference and visualised using SVbyEye v0.99.0^90^. To ensure call accuracy, the UCSC Genome Browser (http://genome.ucsc.edu) was used to examine the repetitive content of candidate complex inversions and HDRs; candidates occupying repeat-rich genomic regions that could confound alignment or SV calling were excluded. Similarly HDR regions were curated and collapsed across samples. This process resulted in a nonredundant set of high confidence variants (see **Supplementary Note 1 and Supplementary Fig. S2-S3**).

#### Tandem repeat genotyping

Tandem repeat (TR) catalogs were generated using the TRACK pipeline^91^ as described in companion manuscript Adam et al 2026. Briefly, TRs were identified using Tandem Repeat Finder v.4.09 ^92^ and filtered based on total repeat length (≤ 10 kb), copy number (≤ 2.5), and constancy score, i.e., percent matches between adjacent copies (≤ 60%). CHM13 coordinates for 66 TRs linked to human expansion disorders were extracted from the STRipy database ^52,92^. These coordinates were lifted to the *Pan troglodytes* assembly using the UCSC Liftover tool^93^. TRs were genotyped separately for both species using Tandem Repeat Genotyping Tools v.3.0 (TRGT;^94^). The merged multisample VCF was filtered for missing data (--max-missing 1), minimum allele spanning depth (>3), and allele constancy score (>=60%), resulting in 61 pathogenic TRs. Mean allele length was computed for both species and compared against the minimum pathogenic length threshold in humans, as reported in STRipy. To test whether heterozygosity differed between pathogenic and nonpathogenic TRs, we used Wilcoxon rank-sum tests.

### Functional effects of SVs and SNVs

Functional consequences for SNVs (biallelic SNPs) and SVs (SVs ≤100 kb; concatenated truvari calls) were predicted using the Ensembl Variant Effect Predictor (VEP) using species-matched references and gene annotations. Variants were categorized into four impact categories: HIGH, MODERATE, LOW, and MODIFIER based on their predicted impact on annotated genes. Site Frequency Spectra (SFS) were generated for each species per impact category. For each species we computed the unfolded SFS of biallelic SNVs and SVs by counting non-reference alleles across diploid long-read genomes (n = 5 bonobos, 24 chimpanzees, 47 humans from the HPRC year-1 release). Fixed sites (all-alt) were excluded. Counts were converted to proportions within each species × impact-classHere, the most severe predicted consequence (HIGH, MODERATE, LOW, MODIFIER) was retained, and impact was binarised as “moderate/high” vs “modifier/low”. For gene-level analyses, we assigned each protein-coding gene the highest impact level observed among its associated variants. Genes were filtered to include only high-confidence protein-coding annotations, excluding uncharacterized LOC symbols and noncoding RNAs (snoRNAs, lncRNAs, and pseudogenes). For downstream constraint analysis, these were simplified into “Moderate/High” and “Modifier/Low” groupings. SV lengths and repeat content (e.g., SINE/Alu, LINE/L1) were summarized to characterize the structural landscape of functional mutations. To ensure comparability across different sample sizes, the proportion of variants at each allele count was calculated. Proportions were visualized using a square-root transformation on the y-axis to facilitate the comparison of rare deleterious alleles versus common neutral variants.

To assess the selective pressures acting on variants of functional impact, we integrated genomic data with Gene Constraint Scores (s_het_) derived from Zeng et al. in humans^50^. Per-gene constraint was taken from the posterior mean s_het_ estimates and genes were classified into four selection regimes based on their posterior mean s_het_. Differences in the distribution of constraint scores between species and impact levels were assessed using two-sided (two-tailed) Wilcoxon Rank-Sum tests to account for the non-normal distribution of s_het_ values. We additionally performed gene Ontology (GO) enrichment analysis to test for biological significance on shared protein-coding genes highly impacted by SVs and SNVs (**Supplementary Fig. S1, Supplementary Table S5-S7**), as well as highly impacted by SVs and SNVs with posterior mean s_het_ matching extreme and strong selection regimes (**Supplementary Table S8-S9**). Analyses were conducted in ShinyGO v.0.85^95^ with a minimum pathway size of 15. Significance was assessed using the false discovery rate (FDR), and only pathways with FDR-corrected p < 0.05 were considered enriched.

### Ancestral recombination graph analyses

SNVs called from long-read haplotype assemblies of chimpanzees and bonobos (this resource) and humans (from both the HPRC year-1 and HGSVC release) mapped to respective T2T references were merged into phased diploid VCFs, restricted to biallelic SNVs, and partitioned into non-overlapping 5 Mb blocks per chromosome, excluding centromeric and telomeric regions. We then reconstructed local genealogies for each species with SINGER, as described in Deng et al^58^. Briefly SINGER was run per 5Mb block per chromosome with default MCMC settings, Ne = 4 × 10^4 for chimpanzees and bonobos, and Ne=2 x 10^4 for humans and μ = 1.25 × 10^-8 per bp per generation. Blocks of the same MCMC index were stitched into chromosome-wide tree sequences with *tskit*, yielding 100 posterior trees per chromosome. We then calculated TMRCA in non-overlapping 1 kb windows as the span-weighted mean root time of marginal trees overlapping the window (**Supplementary Fig. S4**). TMRCA estimates were obtained from *tskit.TreeSequence.diversity* with *branch mode* divided by two. We applied this to all haplotypes per species, and to cohorts of haplotypes unique to each population within species, providing the average pairwise coalescence time in the combined sample (*Tpooled*) and within each population (*Twithin*), as described in Deng et al. All statistics were averaged across the 100 posterior ARGs and converted to millions of years assuming a generation time of 25 years for chimpanzees and bonobos, and 28 years for humans. Population assignments for *Twithin* followed Central, Eastern and Western chimpanzees and human continental regions (genome-wide) and populations (targeted regions). Bonobos were treated as a single population (*Tpooled* = *Twithin*) given limited sample size and lack of population structure. The *Tpooled/Twithin* ratio was computed per 1 kb window per population by dividing the posterior-averaged *Tpooled* by the posterior-averaged *Twithin* of that population. Each 1 kb window was annotated with overlapping genes from the corresponding T2T assembly annotations with bedtools.

Next, we computed the fractional overlap of each 1 kb window with species-specific tandem-repeat catalogues described in companion manuscript Adam et al 2026 and other genome feature annotations from UCSC genome browser for each T2T reference. We then filtered out windows with annotations matching satellites, low-complexity sequence, gaps or rDNA, any centromere-satellite content, and with a tandem-repeat overlap above 5 % and short-read accessibility-mask overlap below 20 %. Sex chromosomes, and hybrid individuals were excluded throughout. All remaining windows with TMRCA older than the human–chimpanzee split (> 6 Mya) were outputted as candidates for long-term balancing selection or trans-species polymorphism. Chimpanzee and bonobo gene annotations were then harmonized to their human orthologues (e.g. PATR-A renamed as HLA-A in chimpanzees) and used to assess shared protein-coding genes with deep coalescence likely maintained by balancing selection (**Supplementary Table S10-S32**). Following Deng et al windows with Tpooled/Twithn above the 99.99th percentile of its per-population genome-wide distribution (**Supplementary Figure S4**) were considered to be signatures of putative selective sweeps that reduce local within-population diversity and were outputted as such (**Supplementary Table S33-S34**). We additionally performed gene Ontology (GO) enrichment analysis to test for biological significance on shared genes with TMRCA predating human-chimp divergence time. Analyses were conducted in ShinyGO v.0.85^95^ with a minimum pathway size of 15. Significance was assessed using the false discovery rate (FDR), and only pathways with FDR-corrected p < 0.05 were considered enriched (**Supplementary Fig. S5**).

### Structural variation analyses of glycophorins and other loci

#### Haplotype structures

Unique human and chimpanzee GYP architectures were identified using PGGB^70,96^ and cosigt^70^ as described in Bolognini et al. All chimpanzee and human (https://github.com/lh3/OpenHGL) haplotypes were annotated as described below. Gene conversion and fusion events were then identified by subdividing genes into 500 bp windows and assigning each window to the identity of the closest matching reference gene. Final clustered structures include all gene configuration / structural haplotypes from PGGB/cosigt pipeline in addition to unique gene fusion and gene conversions identified from tiling window analysis. Haplotypes structures were visualized using SVbyEye^90^. Short-read individuals were genotyped using cosigt^70^ as described in Bolognini et al.

#### Protein structures annotation

All protein isoforms were annotated using Phobius^97^ to identify extracellular domains, intracellular domains, transmembrane domains, and signal peptides.

#### Gene annotation

CAT2 v2.0.0 (https://github.com/ph09/CAT2_smk, an updated version of Fiddes et al 2018^98^) was used to annotate genes on all of the sequences using the CHM13 gene annotations of the GYP genes as the reference gene set. For this, we used the minimap2 genome alignment-based and spliced-transcript alignment based modules to transfer genes from the reference to each of the targets. The AUGUSTUS^99^ v3.5.0 gene prediction module was used to fix gene and CDS boundaries. Gene models for GYPA, GYPB, and GYPE on each of the target sequences were taken from the merged GFF3. For each gene, one representative transcript was chosen: MANE^100^ Select accessions when tagged in the GFF, otherwise the transcript with the most exons. Each target copy was aligned to each reference gene sequence with MAFFT^101^ v7.525 (--auto, global pairwise alignment). Per–query-base identity to the aligned reference column was computed from the alignment; values were smoothed with a 200 bp sliding-window mean along the query. At each position, the reference with highest smoothed identity was taken as the local best match. Contiguous runs of the local winner were merged; runs shorter than 200 bp were absorbed into neighboring runs by mean identity over the short interval. The overall gene label was assigned as the reference whose summed smoothed identity over the full gene was largest. Segments whose local winner differed from that overall label but had a small identity margin over competing references (below 0.02) were reassigned to the overall label. Remaining multi-gene patterns were classified as fusion if any non-primary segment had an identity margin over the next-best reference of at least 0.05, otherwise as gene_conversion; single-gene patterns were normal.

## Data Availability

All raw sequencing data are deposited in NCBI under accession number (*PENDING*). Genome assemblies are deposited under (*PENDING*).

## Code Availability

All code used in the paper can be found in the following GitHub repository https://github.com/sudmantlab/panpan_diversity_project and is archived in zenodo (*PENDING*).

## Acknowledgements

We thank the Sudmant lab, Aida Andres, and Megan Dennis for helpful discussion. This work was supported, in part, by National Institutes of Health (NIH) National Institute of General Medicine award R35GM142916 to PHS, NIH National Human Genome Research Institute award R01HG013017 to PHS and R01HG002385 to EEE, and a Weill Neurohub Award to PHS, EEE, and AP. This manuscript is the result of funding in whole or in part by the NIH. It is subject to the NIH Public Access Policy. Through acceptance of this federal funding, NIH has been given a right to make this manuscript publicly available in PubMed Central upon the Official Date of Publication, as defined by NIH. The content is solely the responsibility of the authors and does not necessarily represent the official views of the National Institutes of Health. E.E.E. is an investigator of the Howard Hughes Medical Institute.

## Author contributions

PHS conceived of the study and experimental design. MWM provided primate cell lines. JLR, AK, and KH performed tissue culture and HMW DNA extraction. EEE and KH provided sequencing. JLR, RNL, CLA, SF, DB, PH, AG, and PHS performed data analysis. YD, NS, BP, EG, AP, RVF, and EEE provided input on analysis. JLR, RNL, SF, CA, and PHS wrote and edited the manuscript with input from all authors. PHS supervised the research.

---

**Extended Data Figure 1.**
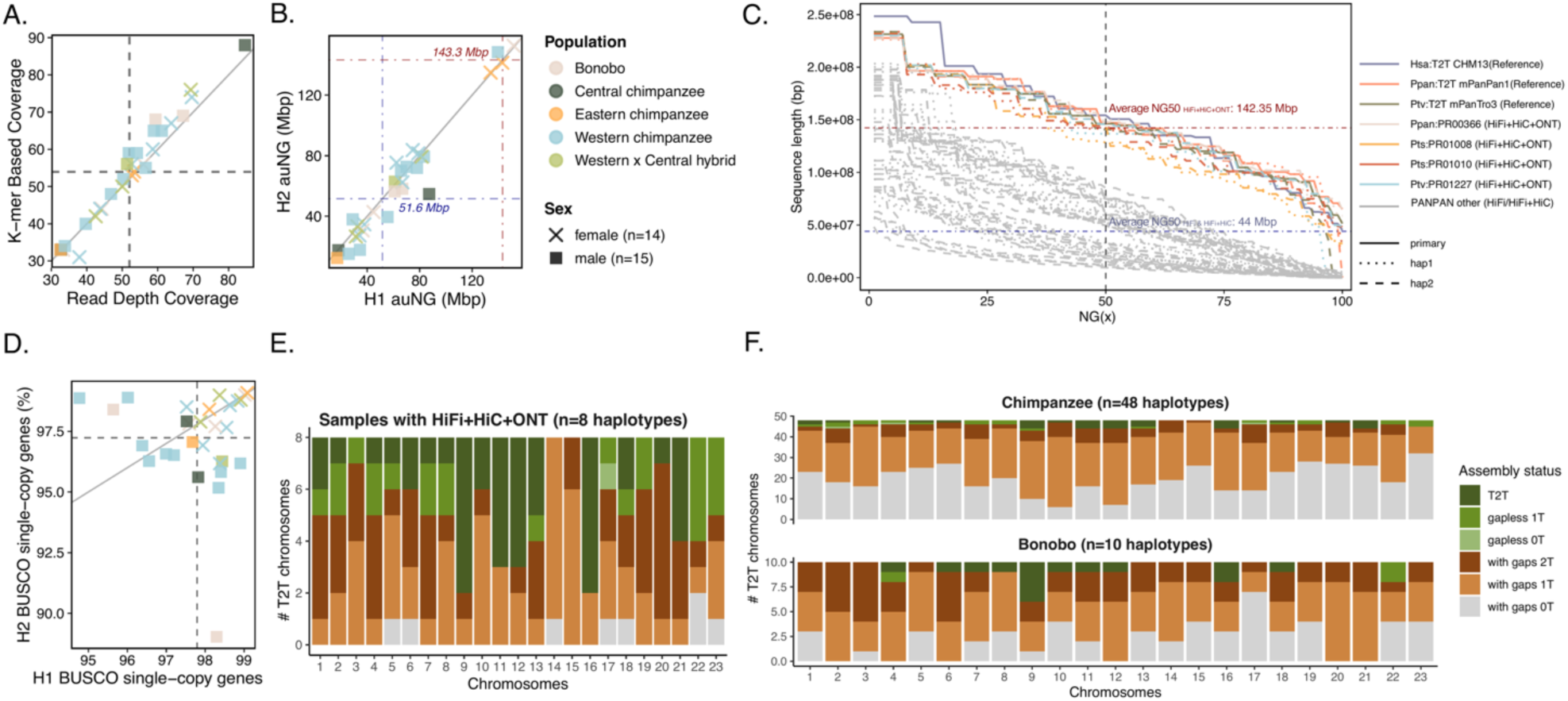
Assembly Quality Control and T2T assembly status. **(A)** Correlation between k-mer based coverage and read depth coverage (average 50x). **(B)** Scatter plot of auNG (area under the NG curve) for Hap1 vs. Hap2, illustrating high consistency between phased haplotypes. **(C)** NG(x) plots showing the distribution of contig lengths across the genome. Solid lines highlight the performance of Verkko assemblies (n=8 haplotypes) relative to T2T human (CHM13 primary assembly) and ape references (chimpanzee mPanTro3 and bonobo mPanPan1 haplotype assemblies). **(D)** Gene completeness as a percentage of BUSCO single-copy orthologs detected in each haplotype from each genome assembly. **(E-F)** Stacked bar plots showing the assembly status for each autosomal chromosome on **(E)** Verkko-assembled haplotypes (n=8) and **(F)** all PANPAN haplotypes. “T2T” signifies a gapless chromosome with telomeres on both ends. Approximately 23.4% of Verkko-assembled chromosomes achieved complete T2T status, 50.5% of chromosomes have both telomeres present (regardless of gaps), and 36.4% of the chromosomes are gapless (regardless of telomeres).

**Extended Data Figure 2.**
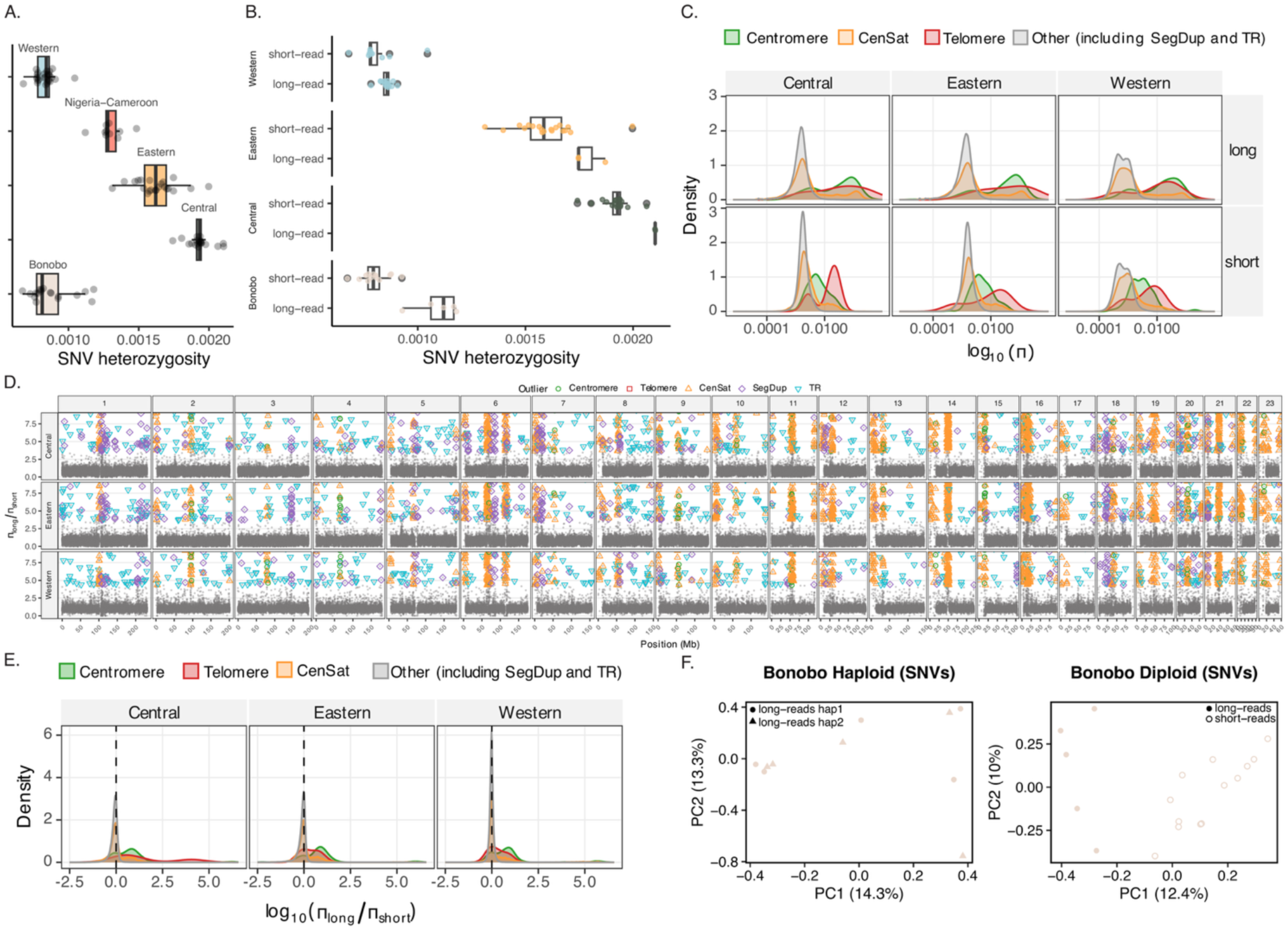
SNV Diversity. **(A)** Genome-wide SNV heterozygosity by population estimated from both Illumina short (n=72) and PacBio HiFi long-read (n=29) diploid individuals, ordered by population and diversity levels. **(B)** SNV heterozygosity estimated from long-read vs. short-read data, per population. Boxplots show the distribution of per-individual Heterozygosity colored by population. Long-read estimates are significantly higher than short-read in every population (Wilcoxon rank-sum, Bonferroni-corrected p < 0.05: Bonobo p = 9.3 × 10⁻⁴, Central p = 0.042, Eastern p = 0.036, Western p = 0.026), and the read-type effect is also significant jointly after controlling for population (linear model, p = 1.1 × 10⁻⁷). **(C)** Density of log₁₀(π_long) and log₁₀(π_short) per 10kb window by read-type and chimpanzee population. Within each panel, the density of per-window π is shown on a log₁₀ x-axis, separately for windows that overlap with genomic features. **(D)** π_long / π_short per 10 kb window across chimpanzee autosomes. Each point is a 10 kb non-overlapping window. Colored markers highlight the top 1% outlier windows annotated with overlapping genome features (Centromere, Telomere, CenSat, SegDup, or TR). The y-axis shows the raw ratio of long-read π to short-read π for that window. Windows where both long and short π were zero were dropped; windows where short π was zero or short reads called no sites were assigned ε = 1 / median(count_comparisons_short) ≈ 1.7 × 10⁻⁷ for the denominator. **(E)** Density of log₁₀(π_long / π_short) per 10 kb autosomal window, faceted by chimpanzee population, colored by feature. Vertical dashed line marks log₁₀(ratio) = 0 (≈ equal long/short π). Density curve shifts indicate that windows have higher π in long-read data than in short-read data. **(C-E)** show that increases of long-read π are concentrated at complex regions and are not a genome-wide effect. **(F)** Principal component analysis (PCA) of bonobo SNV variation mapped to the T2T mPanPan1 reference. Left: haploid PCA of SNVs called from long-read haplotype-resolved assemblies (n = 5 individuals, 10 haplotypes, n= 7,904,333 SNPs after minor allele frequency < 0.05 filter). Right: diploid PCA of SNVs jointly called from read-mapped data, combining long reads from this study (n = 5) with short reads (n=13) from the Great Ape Genetic Diversity Project (total of 18 individuals, n= 7,906,678 biallelic SNPs after minor allele frequency < 0.05 filter). Points are coloured by population; shapes distinguish data type (filled circles, long-read; open circles, short-read; triangles, long-read haplotype 2). Axis labels indicate the percentage of variance explained by PC1 and PC2.

**Extended Data Figure 3.**
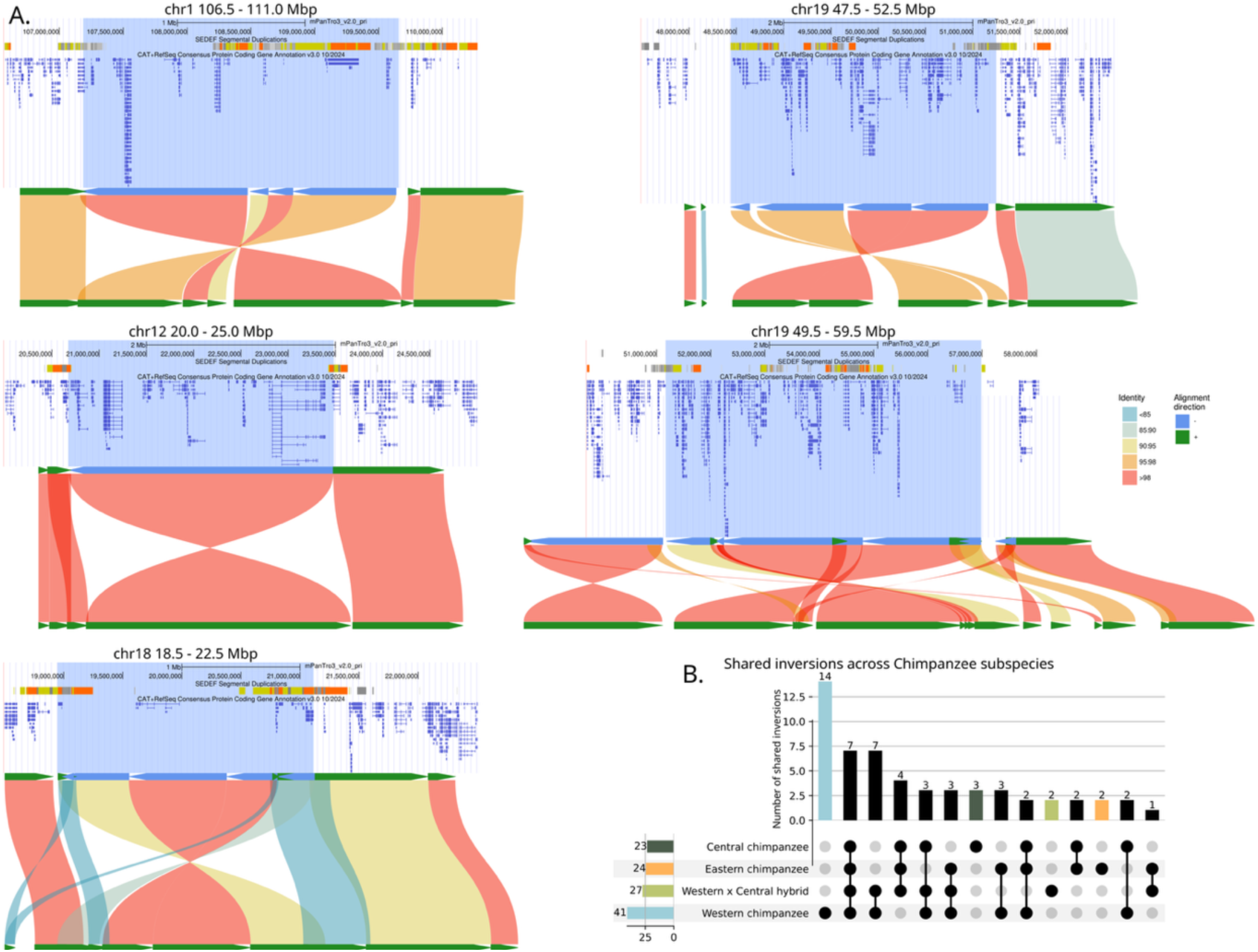
Example genomic architecture and population distribution of structural inversions across chimpanzee subspecies. **(A)** Sequence alignments and genomic context for five representative inversion loci. Each panel shows a single-haplotype alignment (SVbyEye) alongside the corresponding UCSC Genome Browser view spanning the inversion breakpoints. Browser tracks show segmental duplications and genes within and adjacent to each inversion, corresponding to the sequence alignments supporting each call. Coordinates refer to the mPanTro3 primary assembly: chr1: 106.5 - 111.0 Mb, chr12: 20.0 - 25.0 Mb, chr18: 18.5 - 22.5 Mb, chr19: 47.5 - 52.5 Mb, and chr19: 49.5 - 59.5 Mb. **(B)** UpSet plot showing the sharing of inversion polymorphisms across the four chimpanzee subspecies, with intersections reflecting the frequency and overlap of inversions between populations.

**Extended Data Figure 4.**
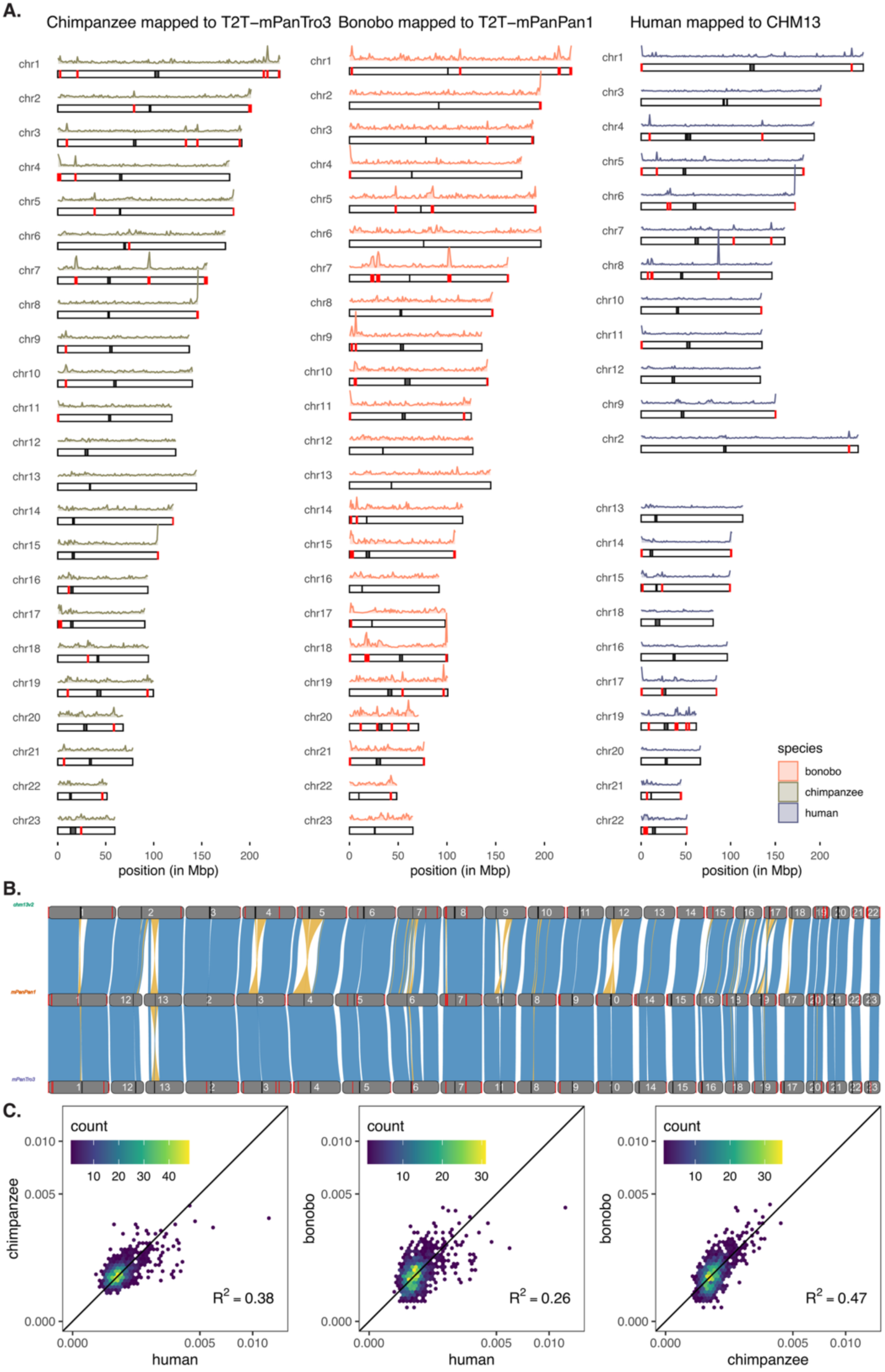
SV Diversity. **(A)** Ideograms of SV density in chimpanzees (left), bonobos (center), and humans (right). Each rectangle represents a chromosome in species-specific reference genomes. Centromere locations are marked with black boxes. Density plots above each chromosome show SV density in non-overlapping 1 Mb windows. SV hotspots are defined as windows with SV density higher than three standard deviations above the mean and are marked by red boxes. The human chromosomes are reordered to maximize correspondence with their homologs in the *Pan* genus. **(B)** A synteny plot of reference genomes in humans (top), bonobos (middle), and chimpanzees (bottom). Synteny between genomes is shown with blue ribbons when in the direct orientation, and yellow when in the reverse orientation. Centromere locations are represented by black lines. SV hotspots are marked by red lines. **(C)** Heatmaps showing correlation between normalized SV density in each pair of species. For humans, SV density is computed in non-overlapping 2Mb windows. Each 2Mb window in the human genome is then lifted over to the chimpanzee and bonobo genomes respectively, and the density of SVs in each lifted region is calculated. Lifted regions shorter than 1Mb or longer than 4Mb in total are filtered out. These window-based SV densities are then normalized in each species so that they add up to 1 and plotted in a heatmap between pairs of species, where colors correspond to the number of windows in each hexagonal grid. A 1:1 line is drawn in each plot for comparison, and the coefficient of determination (R^2^) is shown in each plot. Both axes are shown in square root scales.

**Extended Data Figure 5.**
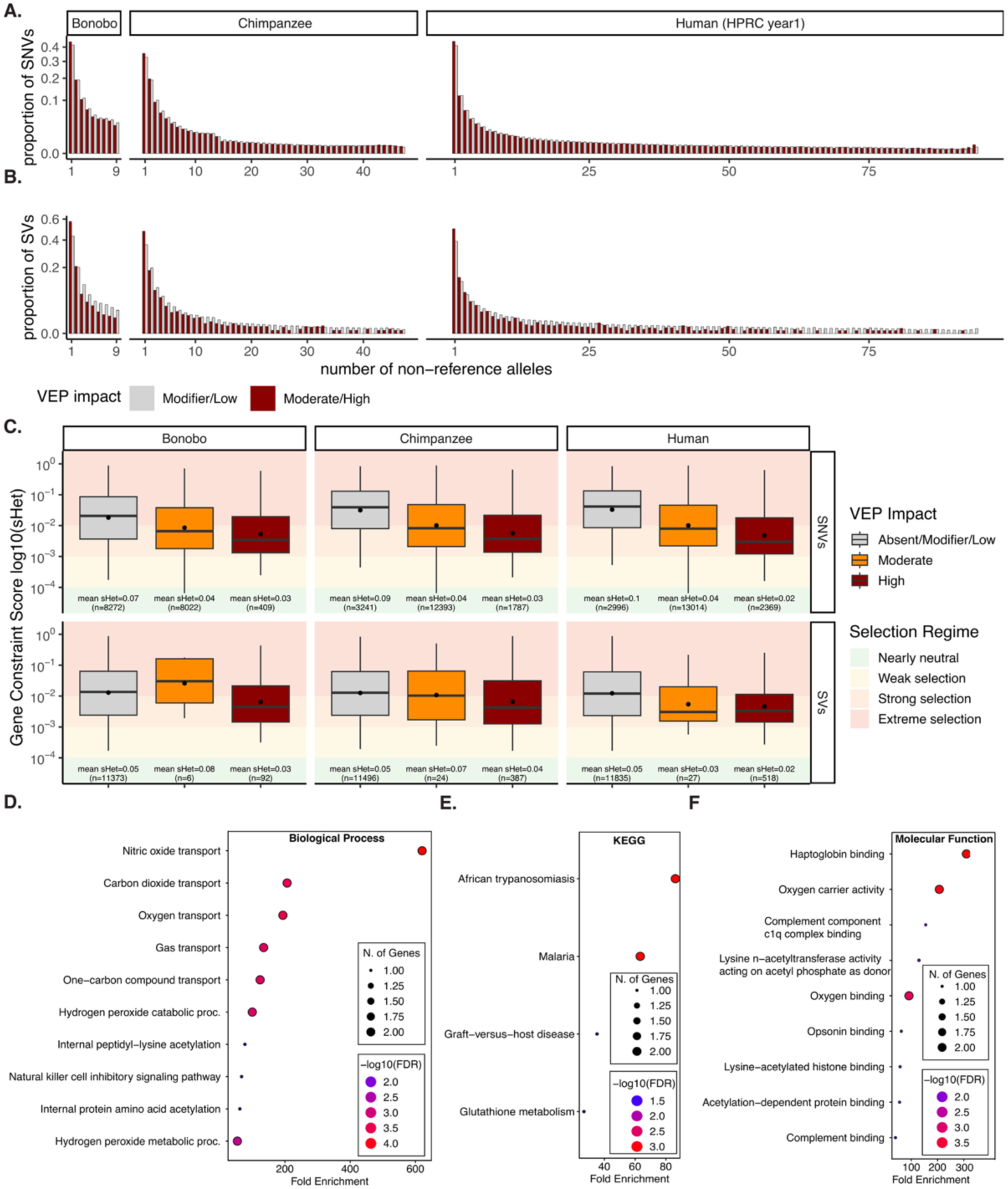
Site Frequency Spectra of genetic variation and gene constraint by variant effect prediction. Site Frequency Spectra (SFS) for SNVs **(A)** and SVs **(B)** for Bonobos, Chimpanzees, and Humans. Bars are colored by Variant Effect Prediction (VEP) impact, comparing “modifier/low” (light grey) to “moderate/high” (dark red) consequences. The y-axis represents the proportion of variants on a square-root scale to highlight differences in rare vs. common variants. **(C)** Distribution of Gene Constraint Scores (s_het_ log10) categorized by VEP impact levels (Absent/Modifier/Low, Moderate, High) for both SNVs and SVs in each species. Boxplots show median and IQR; whiskers extend 1.5 × IQR; black dots mark the arithmetic mean s_het_. Sample size (number of genes) and mean s_het_ are annotated below each box. Background shading represents selection regimes: Nearly Neutral (green), Weak (yellow), Strong (orange), and Extreme (red). High-impact variants are consistently associated with genes under higher evolutionary constraint (lower s_het_). **(D-F)** Gene ontology enrichment analysis of highly constrained genes that are also highly impacted by SVs in both chimpanzees and humans, using different gene ontology sets: **(D)** biological processes, **(E)** KEGG and **(F)** molecular function.

**Extended Data Figure 6.**
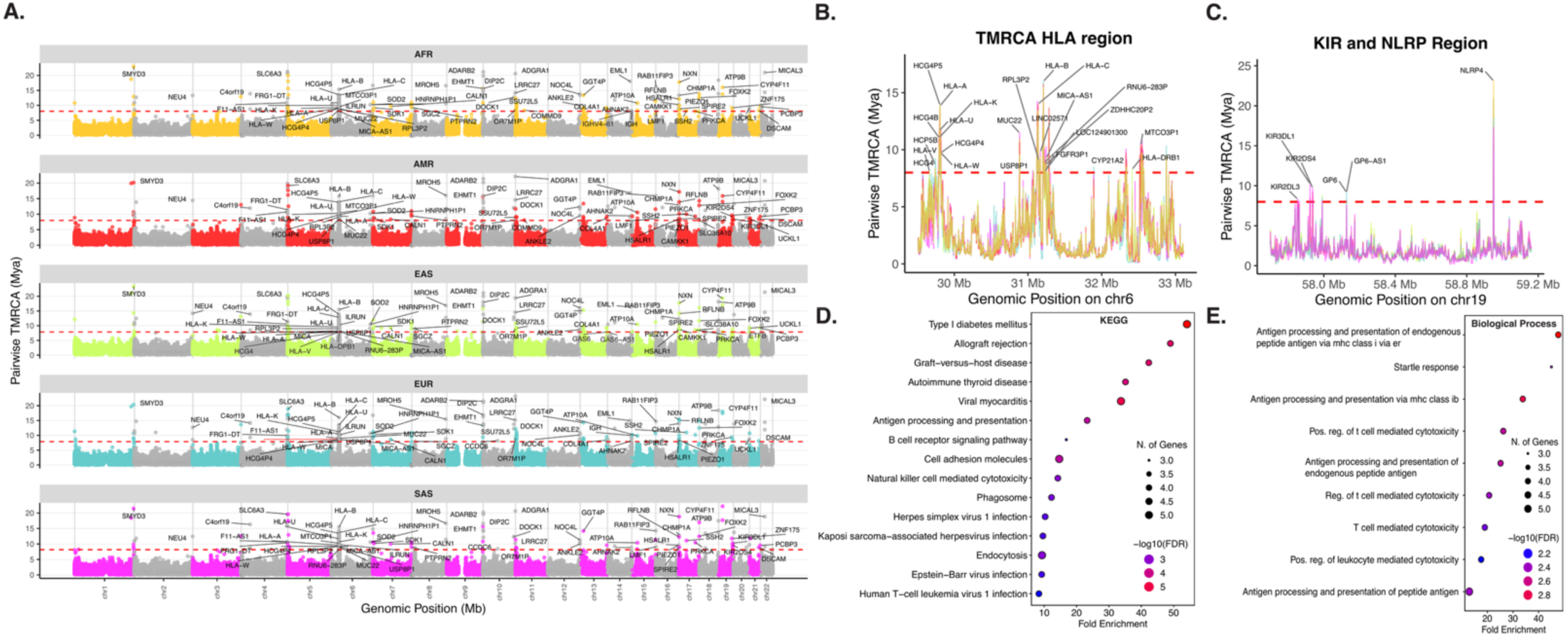
ARG-based inference of deeply coalesced regions in humans. **(A)** Manhattan plots of a genome-wide scan for loci with exceptionally ancient coalescence times in five human continental populations, using average pairwise TMRCA for 1 kb windows. Dashed red lines indicate the 99.99th percentile of the genome-wide empirical distribution. Named genes intersecting with outlier windows are annotated. **(B-C)** Fine-scale examples of HLA **(A)** and KIR/NLRP **(B)** regions, with exceptionally high average pairwise TMRCA in human populations; genes overlapping windows above the 99.99th percentile of the genome-wide empirical distribution are also highlighted. **(D-E)** Gene ontology enrichment analysis for KEGG **(D)** and biological processes **(E)** terms associated with genes overlapping 1kb windows with with average pairwise TMRCA pre-dating *Homo-Pan* divergence (> 6Mya) that are shared between humans, chimpanzees, and bonobos.

**Extended Data Figure 7.**
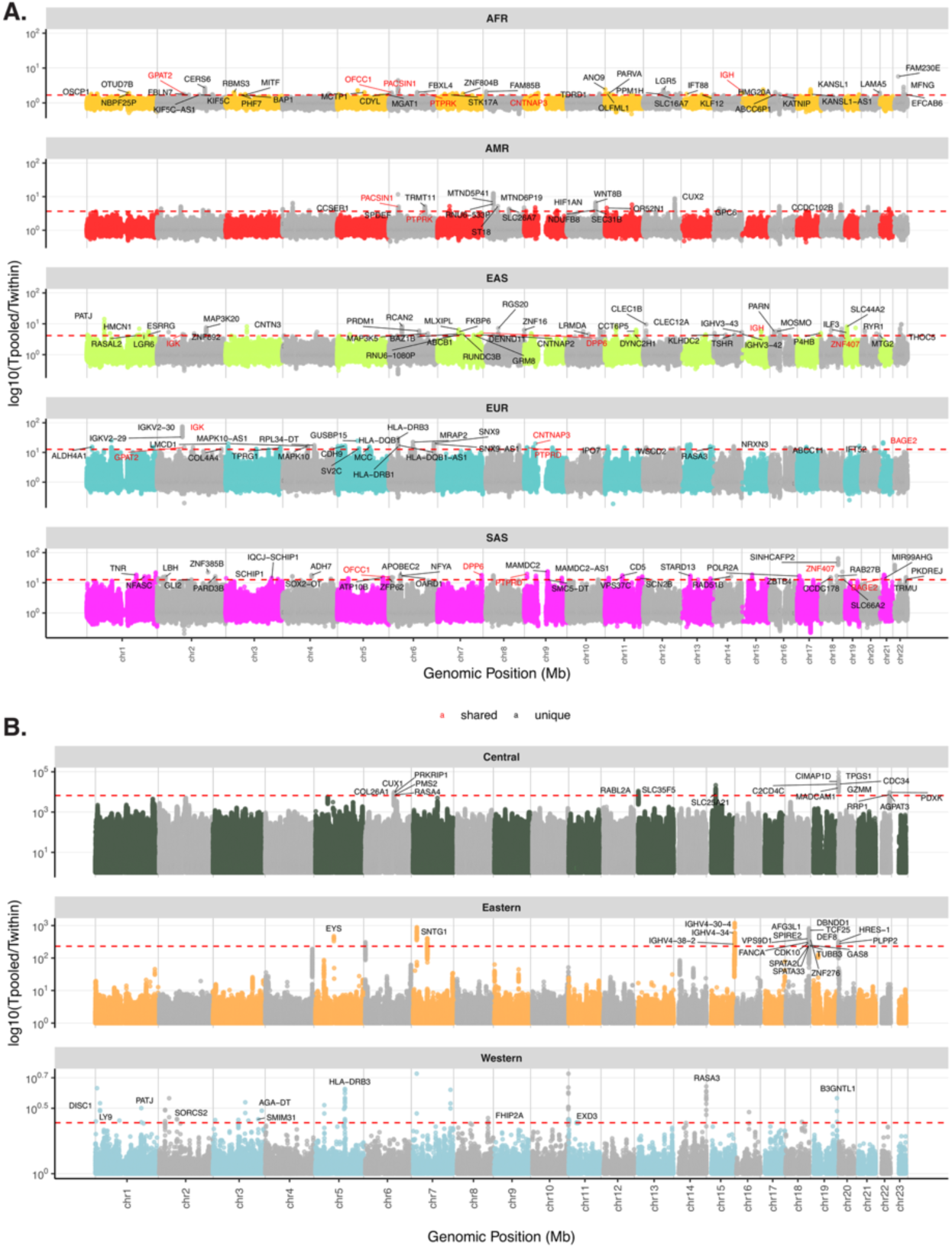
ARG-based detection of population specific reductions in coalescence times in humans and chimpanzees. The log10 of the ratio of the average pairwise coalescence time in the combined sampled (Tpooled) to the average population-specific pairwise coalescence time (Twithin) computed in 1kb windows across the genome in humans **(A)** and chimpanzees **(B)**. Population-specific reductions in local diversity show up as peaks against the genome-wide background. Red dashed lines indicate the genome-wide 99.99th percentile. Named genes intersecting with outlier windows are annotated, and shared cases across human populations are indicated in red.

**Extended Data Figure 8.**
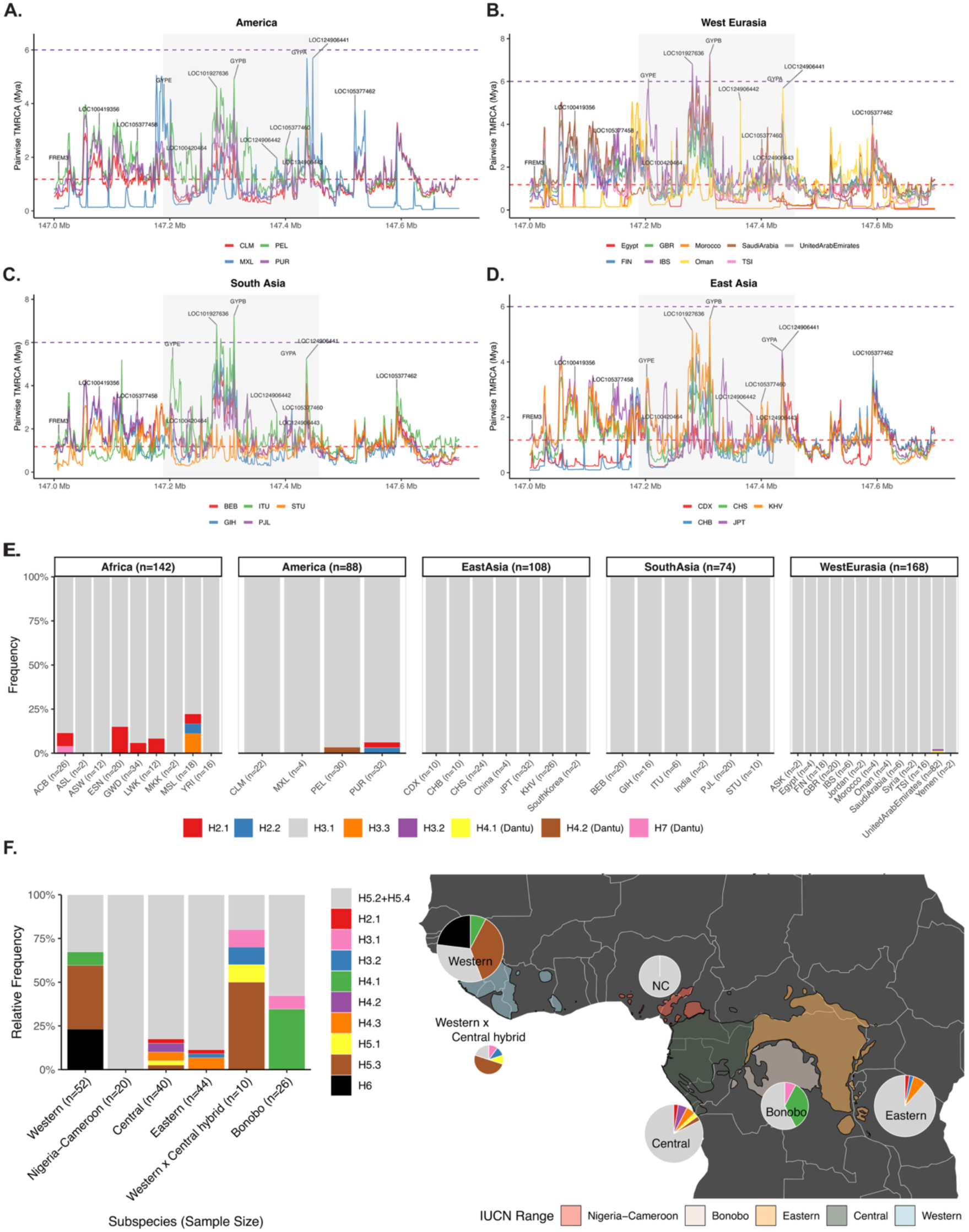
Signatures of deep coalescence at the GYP locus and global distribution of GYP structural diversity in humans and chimpanzees. **(A-D)** Average pairwise TMRCA estimates in 1kb windows across the GYP locus in different human populations, grouped by continent. Dashed lines indicate the genome-wide average TMRCA (red) and the human-chimpanzee split time (purple), with windows overlapping genes highlighted. In each plot, the grey shaded area corresponds to the start and end region of the locus in human T2T reference. **(E)** The frequency of GYP structural haplotypes in different human populations grouped by continent. The number of haplotypes in each population is indicated along the x axis. **(F)** The frequency of GYP structural haplotypes in short-read whole-genome sequencing data of chimpanzee and bonobo populations inferred from haplotype deconvolution, shown in bars (left) and on a map (right). Note that H5.2 and H5.4 cannot be distinguished by this method and are thus combined into a single category.

